# Cognitive hyperplasticity drives insomnia

**DOI:** 10.1101/2024.07.16.603670

**Authors:** Sheng Huang, Chengji Piao, Zhiying Zhao, Christine B. Beuschel, Oriane Turrel, David Toppe, Stephan J. Sigrist

## Abstract

Sleep is vital for maintenance of cognitive functions and lifespan across the animal kingdom. Here, we report our surprising findings that insomniac (*inc*) *Drosophila* short sleep mutants, which lack a crucial adaptor protein for the autism-associated Cullin-3 ubiquitin ligase, exhibited excessive olfactory memory. Through a genetic modifier screen, we find that a mild attenuation of Protein Kinase A (PKA) signaling specifically rescued the sleep and longevity phenotypes of *inc* mutants. Surprisingly, this mild PKA signaling reduction further boosted the excessive memory in *inc* mutants, coupled with further exaggerated mushroom body overgrowth phenotypes. We propose that an intrinsic hyperplasticity scenario genuine to *inc* mutants enhances cognitive functions. Elevating PKA signaling seems to serve as a checkpoint which allows to constrain the excessive memory and mushroom body overgrowth in these animals, albeit at the sacrifice of sleep and longevity. Our data offer a mechanistic explanation for the sleep deficits of *inc* mutants, which challenges traditional views on the relation between sleep and memory, and suggest that behavioral hyperplasticity, e.g., prominent in autistic patients, can provoke sleep deficits.

## Introduction

Sleep is a dynamic process conserved from invertebrates to mammals and humans [1–3]. The molecular, cellular and circuit mechanisms encoding sleep need and executing sleep behavior have been intensively studied [3–5]. Though sleep is proposed to have widespread functions, many studies can be categorized into its restorative functions, which are meant to optimize lifespan and cognition [6]. Indeed, acute- and long-term sleep loss can have adverse effects on both memory and survival [7–10], while efficient sleep benefits against stress, aging and disease [8, 11–13]. Recent studies suggest that multiple tissues and organs mediate the effects of sleep loss, for example gut oxidative stress [8], and fat body metabolism [14, 15]. However, the brain as a signaling hub integrates global information including those from peripheral tissues and circuits for a final behavioral output [16, 17]. The plastic changes in molecular and circuit levels coupling and balancing sleep and memory are thus likely harnessed within core pacemaker neurons and circuits controlling both behavioral processes [18–20]. Indeed, in developmental neurological diseases like autism spectrum disorder, sleep and memory disturbances are prominent [21, 22] and often positively associated with the severity of symptoms in human and animal models [23]. Specifically, sleep deficits and hyper-learning have been associated with brain hyper-connectivity and hyper-plasticity in autistic humans and the valproic acid-induced autism model [24–26]. Indeed, a direct coupling between cellular and behavioral hyperplasticity provides a possible explanation for the etiology of autism spectrum disorder [25]. However, direct causal relations have hardly been established and cellular-genetic models remain scarce.

The fruitfly *Drosophila melanogaster* has long been used to study associative learning and memory [27]. Flies also exhibit sleep states similar to mammals and humans, with reduced locomotion, increased arousal threshold, and compensatory sleep rebound upon sleep loss [28, 29]. Similar to the mammalian hippocampus [30], the mushroom body as a high level integration center of the fly brain plays essential roles in both memory and sleep regulation [19, 20, 31]. Activating or silencing mushroom body intrinsic neurons (Kenyon cells) can change the patterns of sleep and wakefulness [19, 20]. Importantly, bidirectional genetic manipulations of the cyclic adenosine 3’, 5’-monophosphate (cAMP) and Protein Kinase A (PKA) signaling suggest that cAMP/PKA signaling negatively regulates sleep [19, 32]. Interestingly, though cAMP/PKA signaling is *per se* indispensable for associative memory, excessive PKA activity can also suppress memory [33, 34]. Conversely, reducing gene dose of PKA catalytic subunit Dc0 protects from age-associated memory decline in *Drosophila* [33, 34]. In the rodent hippocampus, sleep loss reduces cAMP level and long-term potentiation and impairs cognitive functions [30]. While previous findings define both core cellular and molecular underpinnings in the coupling of memory and sleep regulation, but whether and how the modulations of cAMP/PKA signaling in the fly mushroom body might control the balance between the ability of memory function and the level of sleep remains unclear.

*Drosophila* has been exploited to identify a spectrum of evolutionary conserved genes in regulating sleep [3, 4]. Among them, the *insomniac* (*inc*) locus, encoding an adaptor protein for Cullin-3 E3 ligase-mediated ubiquitination, was identified in two independent genetic screens to be essential for promoting sleep [35, 36]. Inc is evolutionary conserved and its mouse orthologs can functionally restore the sleep of *inc* mutants [37]. Inc has strong expression in the mushroom body, and is essential for proper development of the mushroom body circuit as well as for presynaptic homeostasis plasticity [38, 39]. Furthermore, *inc* mutants are hypersensitive to oxidative stress and short-lived [11, 13, 35]. Interestingly, Cullin-3 lesions have been associated with autism spectrum disorder [40], and pan-neuronal Cullin-3 downregulation has been proposed as a novel *Drosophila* autism model recently [41]. However, the cellular and molecular mechanisms by which Inc modulates Cullin-3 function to regulate sleep and wakefulness remain unknown.

In this study, we firstly demonstrate that *inc* mutants, different from several other sleep mutants we tested in parallel, showed robustly increased scores for olfactory aversive learning and memory, despite their severe sleep deficits. To understand the molecular mechanisms underlying the seemingly uncoupling of sleep and memory in *inc* mutants, we performed a genetic modifier sleep screen and identified the PKA signaling pathway in specifically mediating the sleep deficits of *inc* mutants. We further found that PKA kinase activity was elevated in *inc* mutants, while downregulating PKA signaling by removing a gene copy of *Dc0* (*Dc0*/+) substantially rescued their sleep and longevity phenotypes, but not the oxidative stress response and the mushroom body circuit structural phenotypes. The rescue in sleep and longevity is specific to *inc* mutants, as the same manipulation was not beneficial for both sleep and longevity deficits in other previously established sleep mutants. Surprisingly, the exaggerated memory phenotypes of *inc* mutants were further enhanced by *Dc0* heterozygosity, suggesting that *inc* loss-of-function suppresses sleep through elevated PKA signaling, which in turn counterbalances and limits the excessive ability to form new memories in *inc* mutants. Taken together, our data illustrate how an intrinsic PKA modulation limiting the severity of cognitive hyperplasticity defines a dynamic setpoint for sleep homeostasis. The results might also provide a conceptual frame for neurodevelopmental cognitive disorders such as autism, which are meant to be driven by developmentally-rooted disbalances of neuronal and synaptic processes.

## Results

### *insomniac* (*inc*) short sleep mutants display increased scores for olfactory learning and memory

One major role of sleep is believed to maintain cognitive functions [6], as supported by detrimental consequences of acute and chronic sleep loss, and by beneficial effects of artificially induced sleep [11, 42, 43]. The likely reciprocal relationship between sleep and memory formation is still not well understood. By utilizing the well-established Pavlovian aversive olfactory conditioning [44], we initially screened a spectrum of *Drosophila* long and short sleep mutants, reasoning that chronic sleep modulations by these mutations might allow to shed light on the common molecular mechanisms of coding sleep for cognitive functions.

Interestingly, most tested mutants, regardless of whether they were long or short sleepers, displayed either normal or reduced scores compared to wildtype (*wt*)/control flies in short-term memory (STM) measured immediately after training, or middle-term memory (MTM) measured 3 hours after training (Figures 1A-1K). Specifically, the short sleepers *Shaker minisleep* (*mns*) [45], 1xBRP [11, 46], *homer* [47], *wide awake* (*wake*) [48] and *argus* (*aus*) [49] mutants were largely normal in olfactory associative learning and memory (Figures 1A-1K). *Neurofibromatosis-1* (*nf1*) and dopamine transporter mutant *fmn* were previous shown to regulate olfactory memory [50, 51], and we could recapitulate these findings (Figures 1C, 1D and 1F). Interestingly, the long sleeping mutant *Fbxl4* [52] showed a specific decrease in STM but not MTM (Figures 1D and 1G). Surprisingly, a single severe short sleeping mutant, *insomniac^2^* (*inc^2^*), exhibited higher STM and MTM scores (Figures 1D and 1I). MTM is composed of consolidated anesthesia-resistant memory (ARM) and unconsolidated anesthesia-sensitive memory (ASM), which can be dissected through amnestic cooling [11, 27, 53]. The ARM component was specifically increased in *inc^2^* mutant (Figures 1J and 1K). Due to occasions of unavailability of some of these mutant animals during memory screening, not all memory components were tested for each mutant. Given that the *inc^2^* short sleep mutants surprisingly showed enhanced learning and memory, we decided to focus on dissecting the function of Inc in regulating both sleep and memory.

**Figure 1.**
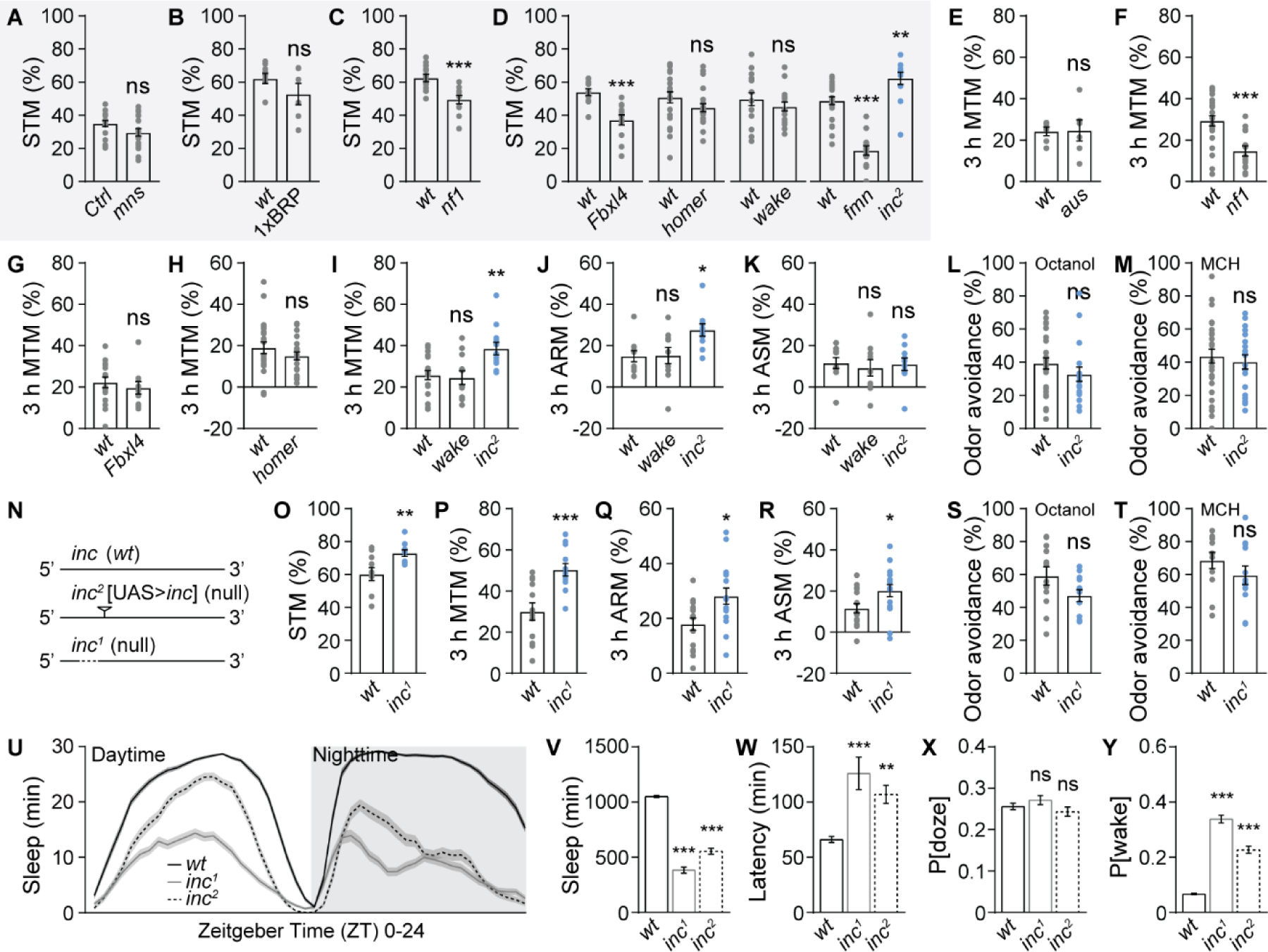
Olfactory learning and memory screen identifies Inc protein as a memory suppressor. (**A-D**) Olfactory short-term memory (STM, or learning) tested immediately after training for *Shaker mns* (**A**), 1xBRP (**B**), *nf1* (***C***), and *Fbxl4*, *homer*, *wake*, *fmn* and *inc^2^* (**D**). n = 18 for *mns*, 6-9 for 1xBRP, 11-12 for *nf1*, 12 for *Fbxl4*, 22 for *homer*, 16 for *wake*, and 12-25 for *fmn* and *inc^2^*. (**E-I**) Olfactory middle-term memory (MTM) tested 3 hours after training for *aus* (**E**), *nf1* (**F**), *Fbxl4* (**G**), *homer* (**H**), and *wake* and *inc^2^* (I). n = 6 for *aus*, 14-22 for *nf1*, 10-16 for *Fbxl4*, 21-23 for *homer*, and 11-19 for *wake* and *inc^2^*. (**J** and **K**) Anesthesia-resistant memory (ARM) (**J**) and anesthesia-sensitive memory (ASM) (**K**) tested 3 hours after training for *wake* and *inc^2^*. n = 10. (**L** and **M**) Odor avoidance of *inc^2^* for octanol (**L**) and 4-methylcyclohexanol (MCH) (**M**). n = 18-29. (**N**) Schematic illustration of both *inc^1^* and *inc^2^* mutants. Note that *inc^2^* is a p-element-mediated mutant with a UAS cassette for driving *inc* expression with the presence of Gal4 [35, 39]. (**O-T**) STM (**O**), MTM (**P**), ARM (**Q**), ASM (**R**) and octanol (**S**) and MCH (**T**) odor avoidance for *inc^1^*. n = 10 for STM, 12 for MTM, 16 for ARM and ASM, and 12 for odor avoidance. (**U-Y**) Sleep structure of *inc^1^* and *inc^2^* flies averaged from measurements over 2-4 days, including sleep profile plotted in 30-min bins (**U**), daily sleep amount (**V**), sleep latency at ZT12 (**W**), P[doze] (**X**) and P[wake] (**Y**). n = 69-96. Except for *mns* mutants, which were in an isogenized *Canton-S* control (*Ctrl*) background, all other mutants were compared to *w^1118^*(*wt*) animals. *p < 0.05; **p < 0.01; ***p < 0.001; ns, not significant. Error bars: mean ± SEM.

*inc^2^* allele is a p-element-mediated *inc* mutation with undetectable Inc protein levels in immunoblots [35, 36]. To further demonstrate the important role of Inc in suppressing olfactory learning and memory, we tested the *inc^1^* null mutant, which harbors a small deletion covering 5’UTR and part of the first exon of *inc* gene (Figure 1N) [35], for STM, MTM, ARM and ASM. Indeed, the *inc^1^* mutant behaved indistinguishable from *inc^2^* mutant in increasing STM, MTM and ARM scores (Figures 1O-1Q), confirming the essential role of Inc in constraining aversive olfactory memories. ASM was also increased in *inc^1^* mutants, potentially reflecting different strengths of these alleles. Importantly, both *inc^1^* and *inc^2^* mutants did not show any obvious alteration in odor sensation, as shown by their normal odor avoidance scores, proving a genuine olfactory memory phenotype in these animals (Figures 1L, 1M, 1S and 1T). Both of the two distinct *inc* mutations were extremely wake-promoting (Figures 1U and 1V) in our hands, fully consistent with previous findings [35, 36]. Interestingly, *inc* mutations provoked sleep loss mainly through an increase of sleep latency and a drastic decrease of sleep quality as depicted by the extremely increased probability of awaking from sleep (P[wake]), but leaving sleep pressure unaffected as indicated by the normal probability of falling asleep from wakefulness (P[doze]) [54]. In short, both *inc* alleles suffer from a substantially decrease of sleep depth and quality (Figures 1X and 1Y). Taken together, these data suggest that *inc* loss-of-function, associated with severely decreased sleep, surprisingly enhances the formation of olfactory memory.

### Genetic modifier sleep screen of *inc* mutants

The conventional concept that sleep is tightly coupled with proper ability of cognitive functions [55] is in contrast with our findings of the *inc* mutants. To better understand this apparent paradox and identify molecular mechanisms in mediating this phenotypic constellation of *inc* mutants, we undertook a modifier screen by testing autosomal heterozygous mutations for a robust modulation of the *inc* sleep phenotypes. We here focused on key players in major signaling pathways and biological processes involved in sleep regulation, circadian rhythms, learning and memory, neurotransmission and synaptic plasticity. We reasoned that any heterozygous hits identified through this screen in interacting with the *inc* mutant sleep phenotypes might allow for insights into the etiology of the *inc* sleep/memory phenotypes.

Loss of *inc* suppresses sleep mainly by promoting nighttime sleep latency (Figure 1W) and decreasing sleep quality indicated by P[wake] (Figure 1Y). Thus, in our modifier sleep screen, we quantified daily sleep amount, nighttime sleep latency and P[wake], and calculated the absolute differences between *inc* mutant flies with or without the presence of heterozygous candidate mutations (Figures 2A-2F). We screened ∼70 isogenized alleles (outcrossed for six generations to *wt*/control background), finding that *inc* mutants were *per se* sensitive for heterozygous modifiers further exaggerating their short sleep phenotypes (Figures 2A and 2D). As we showed previously [46], removing a single copy of the ELKS-family active zone scaffold protein Bruchpilot (1xBRP) was more profound in wake-promoting of *inc* mutants than of *wt*/control background (Figures 2A-2C, 2G and 2H). Inc functions as an adaptor of the E3 ligase Cullin-3 [35]. Consistent with Inc in promoting Cullin-3 function in sleep regulation, *Cullin-3* heterozygosity further enhanced the *inc* sleep deficits (Figures 2A-2C, 2G and 2H). Moreover, genetic modulations of GABAergic signaling did dynamically modify the *inc* sleep phenotypes (Figures 2A-2C, 2I and 2J).

**Figure 2.**
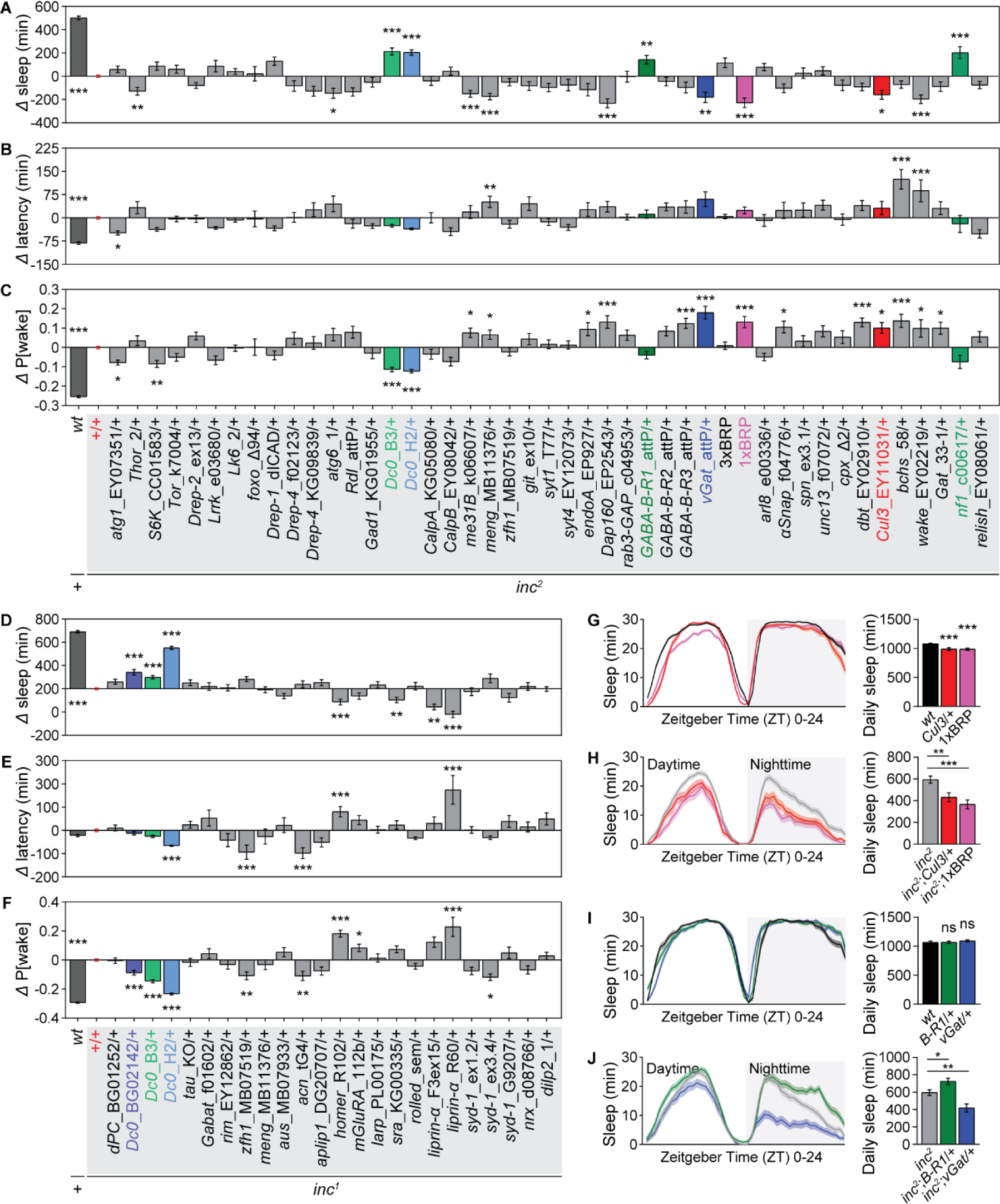
Genetic modifier sleep screen for *inc* mutants. (**A-C**) Genetic screen in *inc^2^* mutant background analyzing the absolute difference in daily sleep (**A**), sleep latency at ZT12 (**B**) and P[wake] (**C**) between *inc^2^* and *wt* or *inc^2^*with autosomal heterozygous candidate mutations. n ≥ 25. (**D-F**) Genetic screen in *inc^1^* mutant background analyzing the absolute difference in daily sleep (**D**), sleep latency at ZT12 (**E**) and P[wake] (**F**) between *inc^1^* and *wt* or *inc^1^*with autosomal heterozygous candidate mutations. Note that some candidates tested in *inc^2^* background were also tested in *inc^1^* background in the screen. n ≥ 12. (**J** and **K**) Some examples of candidates shown to tune sleep of *inc* mutants by their heterozygosity, including *Cullin-3^EY11031^*/+ (*Cul3*/+) and *brp^c04298^*/+ (1xBRP) (**G** and **H**), and *GABA-B-R1^attP^*/+ (*B-R1*/+) and *vGat^attP^*/+ (**I** and **J**). n = 40-111 for **G**, 30-51 for **H**, 24-31 for **I**, and 30-51 for **J**. Statistical comparisons were performed either between *wt* and *inc*, or between *inc* with and without heterozygous candidates. *p < 0.05; **p < 0.01; ***p < 0.001; insignificant comparisons are not shown for simplicity. Error bars: mean ± SEM.

In contrast to these potential hits enhancing the *inc* short sleep phenotypes, we only found a few lines which were able to rescue the *inc* short sleep scenario. The strongest and most robust rescue was observed after establishing heterozygosity for PKA catalytic subunit Dc0, particularly pronounced for the *Dc0^H2^* allele in *inc^1^* background (Figures 2A-2F). Moreover, Neurofibromatosis-1 (Nf1), previously shown to regulate cAMP/PKA signaling in the context of sleep and memory regulation [50, 56], also robustly suppressed the short sleep phenotypes of *inc* mutants by its heterozygosity (Figures 2A-2C). Given the consistent rescue effects after a mild genetic reduction of PKA signaling components, we decided to focus our efforts on the potential relationship between Inc and the cAMP/PKA signaling pathway.

### cAMP/PKA signaling mediates the sleep phenotypes of *inc* mutants

To deepen our understanding towards the exact nature of this rescue scenario, we in detail examined the sleep phenotypes of the two *inc* null mutants in conjunction with heterozygosity of several distinct alleles of the PKA catalytic subunit *Dc0* (Figures 3A-3E). *Dc0^B3^* and *Dc0^H2^* alleles are loss-of-function mutants harboring point mutations within the *Dc0* open reading frame [57]. Heterozygosity of *DcO^B3^* and *Dc0^H2^* were previously shown to trigger moderate but significant reductions in PKA activity [33]. We first found that, while *Dc0^H2^* heterozygosity provoked deeper sleep than *Dc0^B3^* heterozygosity in *wt*/control background indicated by shorter sleep latency and lower P[wake], both alleles did not exhibit major alteration in sleep pattern and daily sleep amount (Figure 3A), indicating that a mild reduction has no considerable sleep disturbance in *wt*/control background.

**Figure 3.**
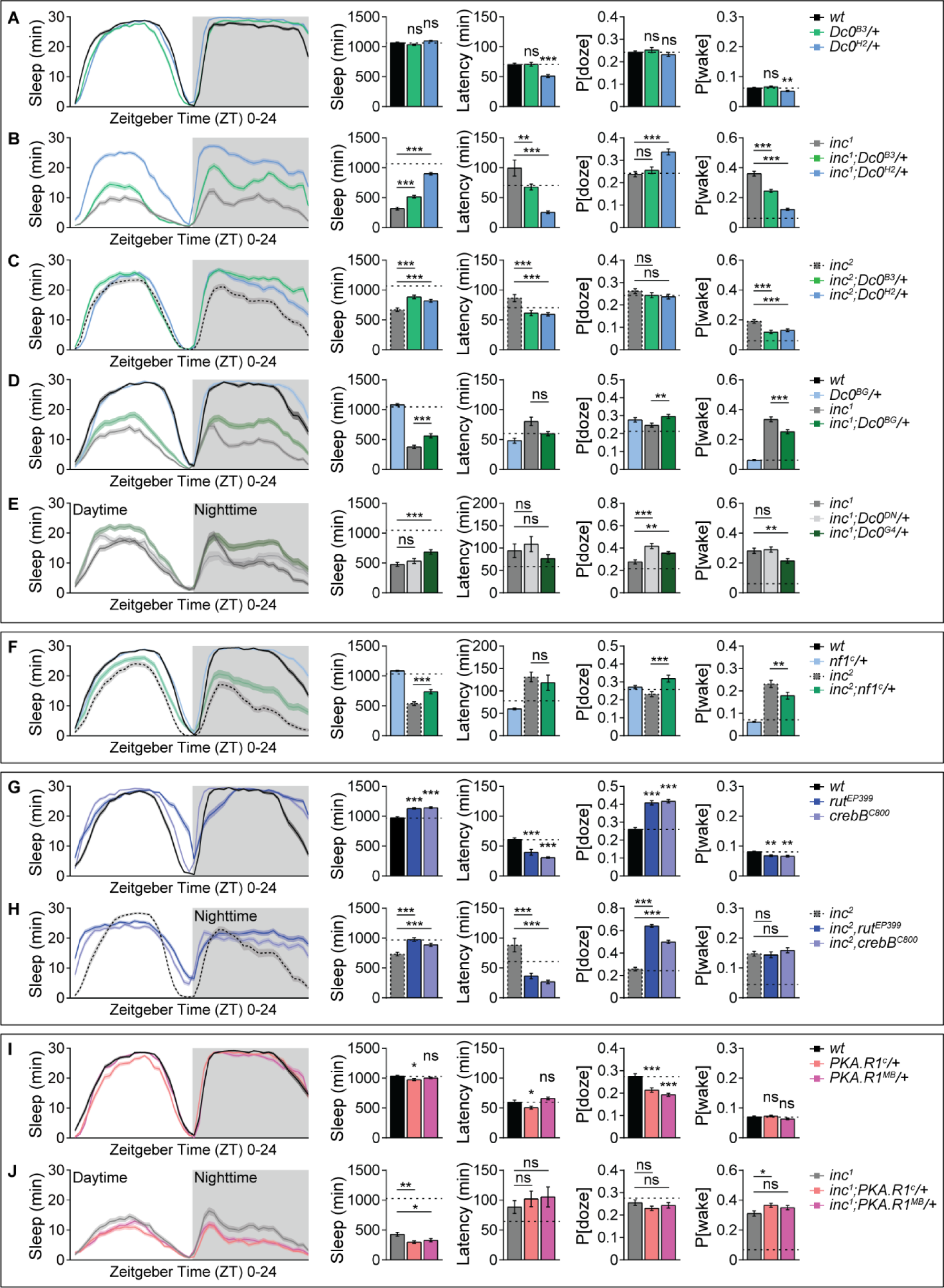
cAMP/PKA signaling mediates the sleep phenotypes of *inc* mutants. (**A**) Sleep structure of *wt*, *Dc0^B3^*/+ and *Dc0^H2^*/+ flies averaged from measurements over 2-4 days, including sleep profile plotted in 30-min bins, daily sleep amount, sleep latency at ZT12, P[doze] and P[wake]. n = 70-76. (**B**) Sleep structure of *inc^1^* with either *Dc0^B3^*/+ or *Dc0^H2^*/+ averaged from measurements over 2-4 days, including sleep profile plotted in 30-min bins, daily sleep amount, sleep latency at ZT12, P[doze] and P[wake]. n = 57-66. (**C**) Sleep structure of *inc^2^* with either *Dc0^B3^*/+ or *Dc0^H2^*/+ averaged from measurements over 2-4 days, including sleep profile plotted in 30-min bins, daily sleep amount, sleep latency at ZT12, P[doze] and P[wake]. n = 73-76. (**D**) Sleep structure of *wt*, *Dc0^BG^*/+ and *inc^1^* with or without *Dc0^BG^*/+ averaged from measurements over 2-4 days, including sleep profile plotted in 30-min bins, daily sleep amount, sleep latency at ZT12, P[doze] and P[wake]. n = 40-71. (**E**) Sleep structure of *wt* and *inc^1^* with either *Dc0^DN^*/+ or *Dc0^G4^*/+ averaged from measurements over 2-4 days, including sleep profile plotted in 30-min bins, daily sleep amount, sleep latency at ZT12, P[doze] and P[wake]. n = 32-48. (**F**) Sleep structure of *wt*, *nf1^c^*/+ and *inc^2^*with or without *nf1^c^*/+ averaged from measurements over 2-4 days, including sleep profile plotted in 30-min bins, daily sleep amount, sleep latency at ZT12, P[doze] and P[wake]. n = 62-80. (**G**) Sleep structure of *wt*, *rut^EP399^* and *crebB^C800^* flies averaged from measurements over 2-4 days, including sleep profile plotted in 30-min bins, daily sleep amount, sleep latency at ZT12, P[doze] and P[wake]. n = 64-70. (**H**) Sleep structure of *inc^2^* with either *rut^EP399^*or *crebB^C800^* homozygous mutants averaged from measurements over 2-4 days, including sleep profile plotted in 30-min bins, daily sleep amount, sleep latency at ZT12, P[doze] and P[wake]. n = 57-62. (**I**) Sleep structure of *wt*, *PKA.R1^c^*/+ and *PKA.R1^MB^*/+ flies averaged from measurements over 4 days, including sleep profile plotted in 30-min bins, daily sleep amount, sleep latency at ZT12, P[doze] and P[wake]. n = 58-63. (**J**) Sleep structure of *inc^1^* with either *PKA.R1^c^*/+ or *PKA.R1^MB^*/+ averaged from measurements over 4 days, including sleep profile plotted in 30-min bins, daily sleep amount, sleep latency at ZT12, P[doze] and P[wake]. n = 61-67. Dashed straight lines indicate mean values of *wt*/control animals tested simultaneously. *p < 0.05; **p < 0.01; ***p < 0.001; ns, not significant. Error bars: mean ± SEM.

Indeed, both daytime and nighttime sleep were substantially restored for both *inc^1^* and *inc^2^* mutants by heterozygosity of either *Dc0^B3^* or *Dc0^H2^* compared to *inc^1^* or *inc^2^* mutants, with shortened sleep latency and drastically reduced P[wake], indicating higher sleep quality (Figures 3B and 3C). As noted earlier (Figure 2D), the sleep restoring effects of *Dc0^H2^* allele were extremely strong in *inc^1^* mutants, further triggering an increase in P[doze], suggesting a higher sleep pressure provoked by this allele (Figure 3B). Interestingly, *Dc0^H2^* heterozygosity was shown to have equal or even slightly stronger PKA activity reduction than *Dc0^B3^* heterozygosity [33]. Thus, the *Dc0* allele-specific sleep effects in *inc* mutant backgrounds might reflect the allelic strength of these PKA alleles.

To further explore the role of PKA signaling in regulating the sleep phenotypes of *inc* mutants and particularly to address whether these effects were *Dc0* allele-specific, we also utilized two p-element-mediated mutants (*Dc0^BG^* and *Dc0^G4^*lines) and one additional dominant negative mutant (*Dc0^DN^* line). These three additional lines are all adult homozygous lethal, but only *Dc0^BG^* and *Dc0^G4^* showed similar effects on restoring sleep to *inc* mutants when compared to *Dc0^B3^* (Figures 3D and 3E), while *Dc0^DN^* as a dominant negative allele distinct from *Dc0* null mutants [58] had not effects on the sleep of *inc* mutants (Figure 3E). Our data thus show that heterozygous *Dc0* null or strong hypomorphic alleles consistently suppress the sleep phenotypes of *inc* mutants, while having little to no effects in *wt*/control animals. In other words, *inc* mutants are sensitized towards changes of PKA signaling, and the extent of PKA signaling seems to be directly relevant for the extent of their sleep deficits.

In inactive state, PKA forms a tetramer consisting of two catalytic and two regulatory subunits (see later Figure 5G). cAMP activates PKA by binding to the regulatory and subsequently releasing the catalytic subunits so as to phosphorylate target proteins. Thus, the regulatory subunit directly antagonizes the kinase function of the catalytic subunit, and the synthesis of cAMP by adenylate cyclases such as Rutabaga (Rut) is essential for activating/disinhibiting PKA signaling. Notably, one of the well-characterized phosphorylation targets of PKA signaling is the cAMP response element binding protein B (CrebB) transcription factor.

Through our genetic modifier sleep screen for *inc* mutants (Figure 2), one additional hit we identified is Neurofibromatosis-1 (Nf1) (Figures 2A-2C). Indeed, *nf1* heterozygosity restored a great extent of sleep in *inc* mutant background by increasing P[doze] and simultaneously reducing P[wake], while it had very mild effects in *wt*/control background (Figure 3F). In *Drosophila*, Nf1 has been intensively studied in the regulation of circadian rhythms [59], sleep [56, 60] and memory (Figures 1C and 1F) [50]. Importantly, Nf1 was shown to be proteasomal-degraded through Cullin-3 E3 ligase-mediated ubiquitination *in vitro* [61]. Furthermore, Nf1 promotes cAMP/PKA signaling pathway by increasing Rutabaga (Rut) adenylate cyclase activity and subsequently the synthesis of cAMP and activation of PKA signaling [50]. Thus, it seems possible that a hierarchical signaling cascade, normally elicited by Inc/Cullin-3 complex-mediated Nf1 degradation and subsequent PKA activity suppression, feeds forward to fine-tune the levels of sleep (See discussion).

In addition, we further found that down-regulating cAMP/PKA signaling by using *rut* and *CrebB* homozygous mutants triggered a significant increase in sleep in both *wt*/control and *inc* mutant backgrounds (Figures 3G and 3H). This increase of sleep was mainly driven by a drastic increase of P[doze] (Figures 3G and 3H). Together with the rescue effects with *nf1* heterozygosity (Figure 3F), our data support that inhibiting PKA signaling by modulating both upstream regulatory components and downstream targets consistently restores sleep in *inc* mutants.

To potentially tune PKA signaling bidirectionally, we further tried to increase PKA signaling by establishing heterozygous scenarios for PKA regulatory subunit type 1 (PKA.R1) both in *wt*/control and *inc* backgrounds. Indeed, opposite to *Dc0* heterozygosity (Figures 3A-3E), the heterozygosity of two alleles of *PKA.R1* exaggerated the short sleep phenotypes of *inc* mutants, while having very mild or no obvious effect in *wt*/control background (Figures 3I and 3J). These data again suggest that *inc* mutants are sensitized towards bidirectional changes of PKA signaling, which seems to directly mediate the sleep phenotypes of *inc* mutants.

### The effects of *Dc0* heterozygosity on promoting sleep are specific to *inc* mutants

The PKA kinase targets a spectrum of downstream effectors and has widespread effects on various biological processes [62, 63]. We next asked whether the effects of *Dc0^H2^* heterozygosity on sleep might be specific for *inc* mutants (Figures 3A-3C, 4A and 4B), or generalize to other genetic sleep deficit scenarios as well. For this purpose, we chose two different approaches: 1) distinct short and long sleep mutants (Figures 4C-4H, 4K and 4L) and 2) activating previously described wake-promoting sleep circuits (Figures 4I and 4J). For most of these scenarios, we did not observe any interference by *Dc0^H2^* heterozygosity on daily sleep, including the short-sleeping voltage-gated potassium channel *Shaker mns* mutants [45] (Figure 4C), *sleepless* (*sss*) mutants [64] (Figure 4D), 1xBRP [11, 46] (Figure 4E), *wide awake* (*wake*) mutants [48] (Figure 4G), *molting defective* (*mld*) mutants [65] (Figure 4H), and long-sleeping *mGluRA* mutants [66] (Figure 4K) and *T-type like voltage-gated calcium channel* (*Ca-α1T*) mutants [67] (Figure 4L). Interestingly, however, the short sleep phenotype of *cacophony^H18^* (*cac^H18^*) [68] was exaggerated (Figure 4F). Similar to most tested sleep mutants (Figures 4C-4H, 4K and 4L), the activation of wake-promoting dopaminergic neurons [69] or helicon cells [70] by expressing low-threshold voltage-gated sodium channel NaChBac [71] (Figures 4I and 4J) was not particularly responsive to the *Dc0 ^H2^*heterozygous scenario. Taken together, our data show that *inc* mutants are specifically sensitized towards a reduction of PKA signaling.

**Figure 4.**
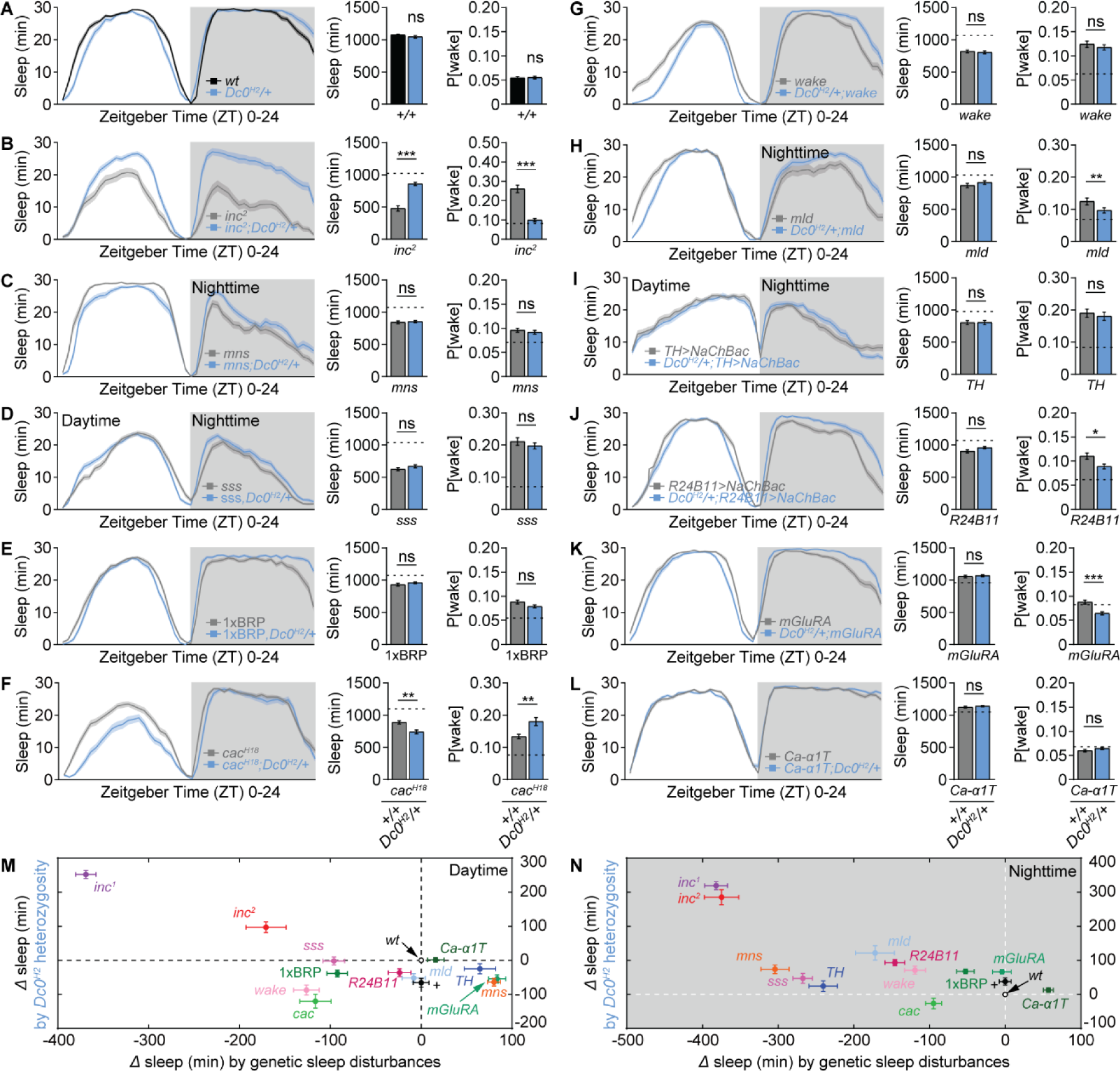
The effects of *Dc0* heterozygosity on promoting sleep are specific to *inc* mutants. (**A**-**L**) Sleep structure of *Dc0^H2^*/+ in control (**A**), *inc^2^* (**B**), *mns* (**C**), *sss* (**D**), 1xBRP (**E**), *cac*H18 (**F**), *wake* (**G**), *mld* (**H**), *TH>NaChBa*c (**I**), *R24B11>NaChBac* (**J**), *mGluRA* (K) and *Ca-α1T* (**L**) backgrounds, averaged from measurements over 2-4 days, including sleep profile plotted in 30-min bins, daily sleep amount, and P[wake]. n = 48-52 for **A**, 32-38 for **B**, 48 for **C**-**F**, 54-55 for **G**, 29-41 for **H**, 47-48 for **I** and **J**, 41-47 for **K**, and 48 for **L**. Dashed lines indicate mean values of *wt*/control animals tested simultaneously. (**M** and **N**) Absolute differences in daytime and nighttime sleep among different genetic sleep disturbances and the diverse effects of *Dc0^H2^*/+ in these backgrounds. *p < 0.05; **p < 0.01; ***p < 0.001; ns, not significant. Error bars: mean ± SEM.

It is worth noting that *Dc0^H2^* heterozygosity drove a slight re-organization of daytime and nighttime sleep, indicated by a slight decrease of sleep during daytime but a mild increase of sleep during nighttime, resulting in an indistinguishable total daily sleep amount (Figures 3A and 4A). Consistently, cAMP/PKA signaling was previously shown to regulate circadian rhythms [72], and *Dc0* expression exhibits daily oscillations [56], suggesting slight different functions for the regulation of daytime versus nighttime sleep. We next systematically compared the effects of *Dc0^H2^* heterozygosity in different backgrounds of sleep disturbance, taking daytime and nighttime into separate considerations (Figures 4M and 4N). Indeed, *Dc0^H2^* heterozygosity was slightly wake-promoting during daytime in most cases, except for *inc* mutants, in which it became sleep-promoting (Figure 4M). During nighttime, *Dc0^H2^* heterozygosity generally showed rather minor sleep-promoting effects, which became much more prominent in *inc* mutant background (Figure 4N).

### Mushroom body PKA kinase activity is increased in *inc* mutants

Inc is broadly expressed in the nervous system [35, 36], including the mushroom body within which Inc was shown to contribute to sleep-promoting effects [39]. Utilizing the *Dc0^BG^* allele, which is a *Gal4* gene trap of *Dc0* locus and its heterozygosity (*Dc0^BG^*/+) exhibited rescue effects on sleep in *inc* mutant background (Figure 3D), to drive GFP reporter expression, we found that Dc0 was also broadly expressed in adult central brain and ventral nerve cord (VNC), with prominent expression in the mushroom body (Figure 5A), consistent with previous reports [33, 73]. To directly monitor the kinase activity levels of PKA, we expressed PKA activity reporter PKA.SPARK [74] in the mushroom body Kenyon cells driven by the mushroom body-specific *VT30559-Gal4* (Figure 5B). Upon PKA activation, SPARK signal forms puncta from otherwise diffused signal *in vitro* and *in vivo* [74, 75]. We indeed observed an increase in the number of PKA SPARK puncta in *inc* mutants, while the average level of SPARK intensity did not change (Figures 5C-5E). We also observed abnormal and overgrown mushroom body morphology in *inc* mutants (Figures 5C and 5F), confirming previous findings [39]. These data suggest elevated PKA kinase activity in *inc* mutant Kenyon cells, which might be directly causal for their severe sleep defects.

**Figure 5.**
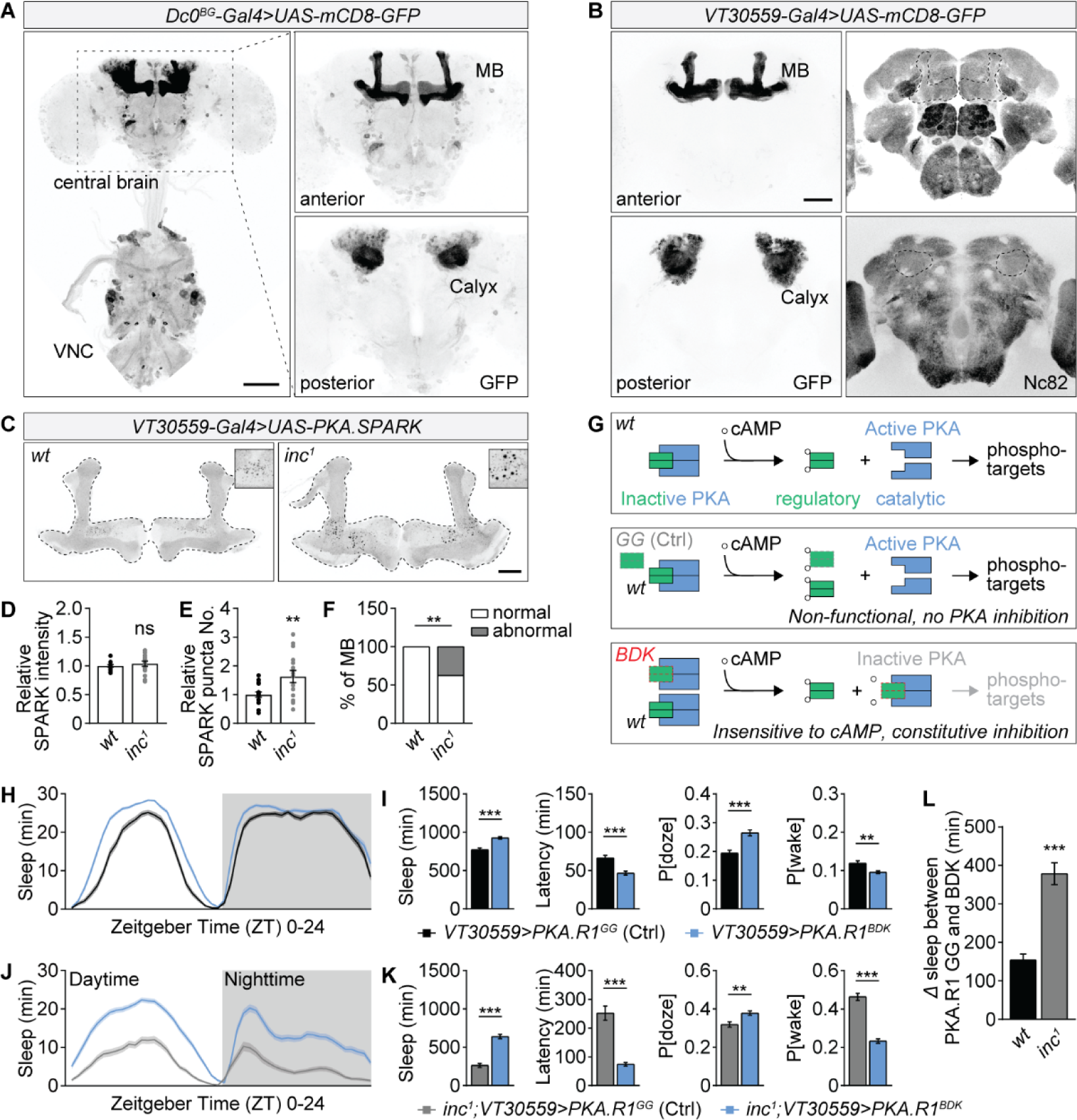
Elevated PKA kinase activity in the mushroom body drives sleep deficits of *inc* mutants. (**A**) Whole-mount brain expressing mCD8-GFP driven by the *Dc0^BG^*Gal4 gene trap line demonstrating the expression pattern of Dc0 in brain and ventral nerve cord (VNC). Dc0 has broad expression but enriched in the mushroom body (MB) lobes and calyx. Native GFP fluorescence is shown. Scale bar: 100 μm. (**B**) Whole-mount brain expressing mCD8-GFP driven the transgenic *VT30559-Gal4* line demonstrating its highly restricted MB-specific expression pattern. The brains were stained with Nc82 antibody, and native GFP fluorescence is shown. Scale bar: 50 μm. (**C-F**) Whole-mount brain immunostaining against GFP for *wt*/control and *inc^1^* flies expressing PKA kinase activity sensor PKA.SPARK in the mushroom body driven by *VT30559-Gal4*, including representative images (**C**), average intensity (**D**) and SPARK puncta number (**E**). *inc* mutants exhibited abnormal mushroom body lobe morphology (**F**). n = 15-16. Scale bar: 20 μm. (**G**) Simplified schematic illustration for the regulation of PKA kinase activity by cAMP and PKA regulatory subunit in *wt* background or under the expression of either a non-functional regulatory subunit (PKA.R1^GG^), which does no longer bind to the catalytic subunit, or a constitutive active regulatory subunit (PKA.R1^BDK^), which constitutively binds to kinase subunit and is insensitive to cAMP. (**H** and **I**) Sleep structure from measurements over 4 days of flies expressing non-functional (GG) or active (BDK) PKA regulatory subunit in the mushroom body driven by *VT30559-Gal4* in *wt*/control background, including sleep profile plotted in 30-min bins (**H**), daily sleep amount, sleep latency at ZT12, P[doze] and P[wake] (**I**). n = 76. (**J** and **K**) Sleep structure from measurements over 4 days of expressing non-functional (GG) or active (BDK) PKA regulatory subunit PKA.R1 in the mushroom body driven by *VT30559-Gal4* in *inc^1^* mutant background, including sleep profile plotted in 30-min bins (**J**), daily sleep amount, sleep latency at ZT12, P[doze] and P[wake] (**K**). n = 83-85. (**L**) Absolutely difference in daily sleep between flies expressing non-functional (GG) and active (BDK) PKA regulatory subunit in the mushroom body in *wt*/control or *inc^1^* mutant background. n = 76-83. **p < 0.01; ***p < 0.001; ns, not significant. Error bars: mean ± SEM.

As mentioned above already, PKA catalytic subunit kinase activity is suppressed by its regulatory subunit [76] (Figure 5G). cAMP binds to the regulatory subunit and disinhibits PKA catalytic subunit (Figure 5G). As shown above, reducing the level of PKA regulatory subunit could further reduce the sleep of *inc* mutants (Figure 3J). To further demonstrate whether PKA kinase activity within the mushroom body is causal for the sleep deficits of *inc* mutants, we expressed an active form of the regulatory subunit (PKA.R1^BDK^), which is insensitive to cAMP and constitutively inhibits the catalytic subunit (Figure 5G) [77], to suppress PKA kinase activity in the mushroom body for sleep measurement. As control, a non-functional form (PKA.R1^GG^), which has no binding activity to the catalytic subunit, was expressed in the mushroom body. We found that expression of the active PKA.R1 in the mushroom body driven by *VT30559-Gal4* triggered a moderate increase of sleep compared to the non-functional form in control background (Figures 5H and 5I). Importantly, mushroom body expression of active PKA.R1 was extremely sleep-promoting in *inc* mutant background (Figures 5J-5L), similar to the effects of *Dc0^H2^* heterozygosity (Figures 3B, 3C, 4A and 4B). These data support that a genuine higher PKA kinase activity within the mushroom body contributes to the *inc* sleep phenotypes.

### Elevated PKA signaling contributes to the compromised life expectancy of *inc* mutants

Similar to other short sleep mutants, *inc* animals were previously shown to be short-lived [35] and hypersensitive to oxidative stress [13]. Given the strong rescue effects of *Dc0* heterozygosity toward the *inc* mutant sleep deficits, we hypothesized that it might also rescue their survival under oxidative stress and longevity phenotypes. For this purpose, we first measured the survival of the *Dc0^H2^* heterozygous animals treated with 2% hydrogen peroxide (H_2_O_2_) and their lifespan under normal aging. Interestingly, *Dc0^H2^* heterozygous animals were more vulnerable than *wt*/control in response to 2% H_2_O_2_ treatment (Figure 6A). In *wt*/control background, the lifespan of *Dc0^H2^* heterozygous animals were significantly increased compared to *wt*/control (Figure 6B), consistently with a previously reported tendence of increased lifespan in *Dc0^H2^*heterozygous flies [33]. We then tested *Dc0^H2^*heterozygosity in both *inc^1^* and *inc^2^* mutants. Similar to a previous report [13], we confirmed that *inc* mutants were hypersensitive to H_2_O_2_ treatment, while *Dc0^H2^*heterozygosity did not have any additive effect here (Figures 6C and 6E). However, in terms of longevity, *inc^1^* mutants were clearly improved and *inc^2^* mutants were fully rescued to the extent of *Dc0^H2^* heterozygous animals (Figures 6D and 6F). These improvements in longevity, together with a strong rescue in sleep (Figures 3B and 3D), suggest a healthier state being established for *inc* mutants by a mild down-regulation of PKA signaling. Again, to test the specificity of this rescue of longevity in the absence of a rescue in the H_2_O_2_ response, we wondered if *Dc0^H2^* heterozygosity was also sufficient to improve the survival of other short-sleeping scenarios including *Shaker mns*, *sss* and 1xBRP in H_2_O_2_ response and longevity. Notably, however, for these short sleep mutants, no rescue was observed (Figures 6G-6L). Thus, our data suggest the scenario that a moderate downregulation of PKA signaling is sufficient to specifically promote longevity as a trait specific to *inc* among the tested short sleep mutants.

**Figure 6.**
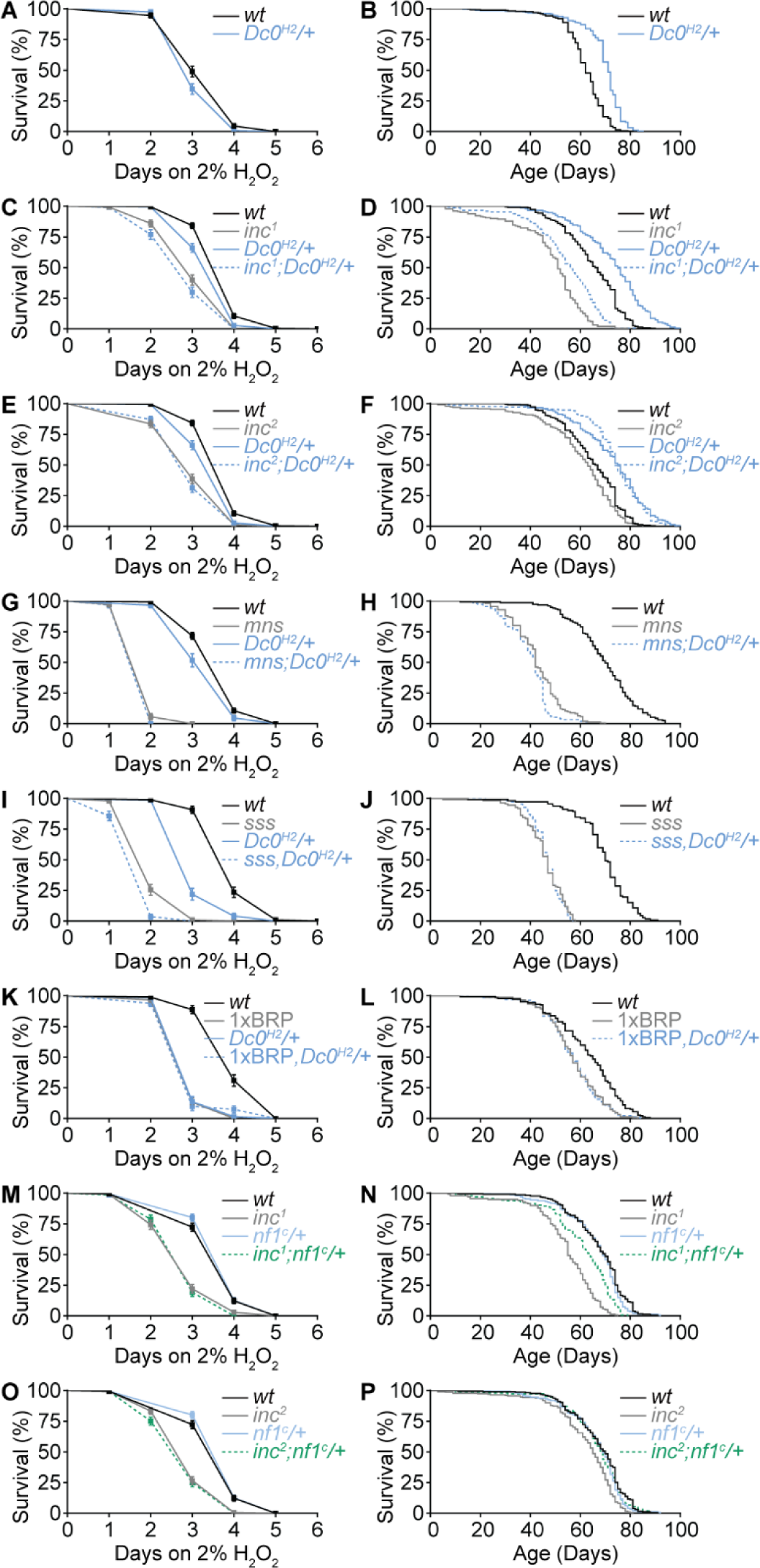
cAMP/PKA signaling mediates the longevity phenotypes of *inc* mutants. (**A** and **B**) Survival curves under 2% H_2_O_2_ treatment (**A**) or normal aging (**B**) for *wt* and *Dc0^H2^*/+ flies. For H_2_O_2_ treatment, n = 133 for *wt* compared to *Dc0^H2^*/+ (n = 124, *p* = 0.025). For longevity, n = 299 for *wt* compared to *Dc0^H2^*/+ (n = 280, *p* < 0.001). (**C** and **D**) Survival curves under 2% H_2_O_2_ treatment (**C**) or normal aging (**D**) for *wt*, *Dc0^H2^*/+ and *inc^1^* with or without *Dc0^H2^*/+. (**E** and **F**) Survival curves under 2% H_2_O_2_ treatment (**E**) or normal aging (**F**) for *wt*, *Dc0^H2^*/+ and *inc^2^* with or without *Dc0^H2^*/+. Note that **C** and **E** share the same *wt* and *Dc0^H2^*/+, as well as **D** and **F**. For H_2_O_2_ treatment, n = 228 for *wt* compared to *Dc0^H2^*/+ (n = 177, *p* < 0.001) or compared to *inc^1^* (n = 143, *p* < 0.001) or compared to *inc^2^* (n = 163, *p* < 0.001), *p* = 0.04 for comparison between *inc^1^* and *inc^1^*;*Dc0^H2^*/+ (n = 117), while no significance (ns) for the comparison between *inc^2^* and *inc^2^*;*Dc0^H2^*/+ (n = 170). For longevity, n = 234 for *wt* compared to *Dc0^H2^*/+ (n = 150, *p* < 0.001) or compared to *inc^1^* (n = 182, *p* < 0.001) or compared to *inc^2^* (n = 232, *p* < 0.001), *p* < 0.001 for comparison between *inc^1^* and *inc^1^*;*Dc0^H2^*/+ (n = 149), and *p* < 0.001 for the comparison between *inc^2^* and *inc^2^*;*Dc0^H2^*/+ (n = 225). (**G** and **H**) Survival curves under 2% H_2_O_2_ treatment (**G**) or normal aging (**H**) for *wt*, *Dc0^H2^*/+ and *mns* with or without *Dc0^H2^*/+. For H_2_O_2_ treatment, n = 264 for *wt* compared to *Dc0^H2^*/+ (n = 87, *p* < 0.001) or compared to *mns* (n = 71, *p* < 0.001), no significance (ns) for the comparison between *mns* and *mns*;*Dc0^H2^*/+ (n = 62). For longevity, n = 156 for *wt* compared to *mns* (n = 156, *p* < 0.001), *p* = 0.001 for the comparison between *mns* and *mns*;*Dc0^H2^*/+ (n = 154). (**I** and **J**) Survival curves under 2% H_2_O_2_ treatment (**I**) or normal aging (**J**) for *wt*, *Dc0^H2^*/+ and *sss* with or without *Dc0^H2^*/+. For H_2_O_2_ treatment, n = 98 for *wt* compared to *Dc0^H2^*/+ (n = 73, *p* < 0.001) or compared to *sss* (n = 98, *p* < 0.001), and *p* < 0.001 for comparison between *sss* and *sss*,*Dc0^H2^*/+ (n = 84). For longevity, n = 110 for *sss* compared to *wt* (n = 112, *p* < 0.001) or compared to *sss*,*Dc0^H2^*/+ (n = 109, ns). (**K** and **L**) Survival curves under 2% H_2_O_2_ treatment (**K**) or normal aging (**L**) for *wt*, *Dc0^H2^*/+ and 1xBRP with or without *Dc0^H2^*/+. For H_2_O_2_ treatment, n = 97 for *wt* compared to *Dc0^H2^*/+ (n = 66, *p* < 0.001) or compared to 1xBRP (n = 91, *p* < 0.001), while no significance (ns) for the comparison between 1xBRP and 1xBRP,*Dc0^H2^*/+ (n = 81). For longevity, n = 150 for 1xBRP compared to *wt* (n = 147, *p* < 0.001) or compared to 1xBRP,*Dc0^H2^*/+ (n = 139, ns). (**M** and **N**) Survival curves under 2% H_2_O_2_ treatment (**M**) or normal aging (**N**) for *wt*, *nf1^c^*/+ and *inc^1^* with or without *nf1^c^*/+. (**O** and **P**) Survival curves under 2% H_2_O_2_ treatment (**O**) or normal aging (**P**) for *wt*, *nf1^c^*/+ and *inc^2^* with or without *nf1^c^*/+. Note that **M** and **O** share the same *wt* and *nf1^c^*/+, as well as **N** and **P**. For H_2_O_2_ treatment, n = 188 for *wt* compared to *nf1^c^*/+ (n = 191, ns) or compared to *inc^1^*(n = 136, *p* < 0.001) or compared to *inc^2^* (n = 180, *p* < 0.001), no significance (ns) for comparison between *inc^1^* and *inc^1^*; *nf1^c^*/+ (n = 141), meanwhile there is also no significance (ns) for the comparison between *inc^2^* and *inc^2^*; *nf1^c^*/+ (n = 177). For longevity, n = 227 for *wt* compared to *nf1^c^*/+ (n = 148, ns) or compared to *inc^1^* (n = 140, *p* < 0.001) or compared to *inc^2^* (n = 232, *p* < 0.001), *p* < 0.001 for comparison between *inc^1^* and *inc^1^*; *nf1^c^*/+ (n = 143), and *p* < 0.001 for the comparison between *inc^2^* and *inc^2^*; *nf1^c^*/+ (n = 148). ns, not significant. Error bars: mean ± SEM.

To test if the rescue effects on longevity of *inc* mutants are specific to *Dc0* heterozygosity or would generalize to other PKA signaling components, we further measured the longevity as well as H_2_O_2_ response of *nf1* heterozygosity in *wt*/control and *inc* mutant backgrounds, which was shown to restore sleep to *inc* mutants (Figure 3F). Interestingly, *nf1* heterozygosity did not show any obvious difference in both survival paradigms compared to *wt*/control (Figures 6M-6P), but substantially improved the longevity of *inc* mutants (Figures 6N and 6P), leaving their H_2_O_2_ response phenotype unaffected (Figures 6M and 6O). Thus, a mild reduction in PKA signaling provoked by the heterozygosity of PKA regulatory component *nf1* triggers similar effects in longevity of *inc* mutants compared to *Dc0* heterozygosity. Taken together, these survival data further support a possible role of Nf1 in molecular signaling transduction from Inc to PKA.

### Intrinsic PKA activity elevation limits the cognitive hyperplasticity of *inc* mutants

Our finding that the intrinsic elevation of PKA signaling underlines the sleep and longevity phenotypes of *inc* mutants triggers the expectation that the boosted memory of *inc* mutants (Figures 1D, 1I-1K and 1O-1Q) might also be moderated by *Dc0* heterozygosity. To directly test this expectation, we first confirmed again the *Dc0^H2^* heterozygosity rescue effects concerning sleep in the same fly cohort measured for middle-term memory (Figure 7A). Surprisingly, though *Dc0^H2^* heterozygosity did not exhibit any significant effect in *wt*/control background, it substantially and significantly exaggerated the excessive 3 h MTM of *inc^2^* mutants (Figure 7B). We further demonstrate that the ARM component was further enhanced by establishing *Dc0^H2^* heterozygosity in *inc^2^*mutants (Figure 7C). Notably, *Dc0^H2^* heterozygous animals displayed re-organized ARM and ASM (Figures 7C and 7D), consistent with a previous report [78].

**Figure 7.**
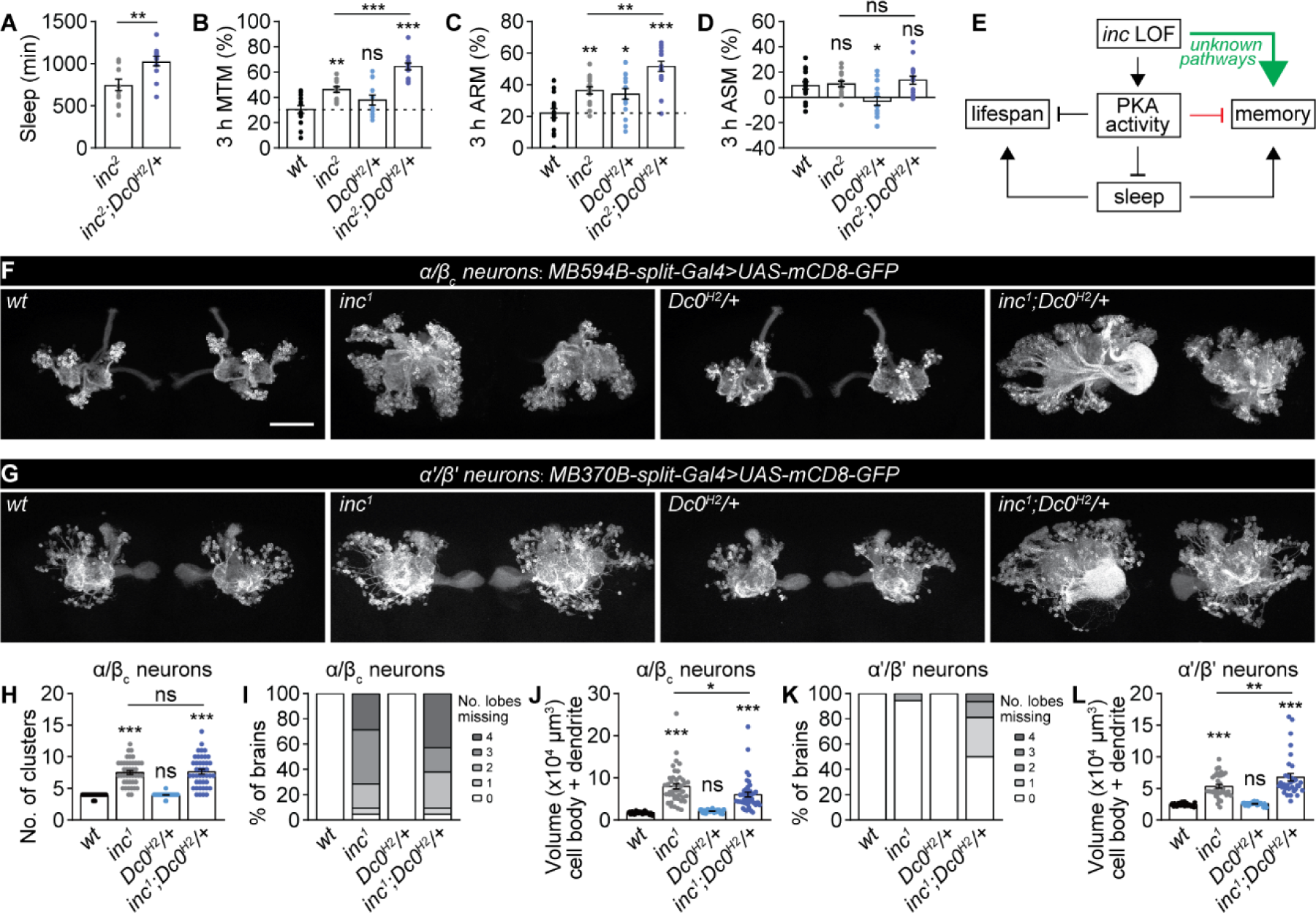
Elevated PKA signaling constrains the excessive memory of *inc* mutants. (**A**) Daily sleep amount of *inc^2^* with or without *Dc0^H2^*/+. *inc^2^*;*Dc0^H2^*/+ animals here were created by genetic combination for olfactory memory, while other identical results were achieved by single round of crosses (Also see methods). (**B-D**) Olfactory memory tested 3 hours after training for *wt*, *Dc0^H2^*/+ and *inc^2^* with or without *Dc0^H2^*/+, including MTM (**B**), ARM (**C**) and ASM (**D**). n = 12-14 for MTM, and 16 for ARM and ASM. (**E**) A simplified model for roles of PKA signaling in coordinating memory, sleep and longevity. *inc* loss-of-function (LOF) is extremely memory-promoting potentially through unknown signaling pathways, which is limited simultaneously by excessive PKA kinase activity. As a consequence, excessive PKA kinase activity provokes severe sleep deficits and compromised lifespan in *inc* mutants. The rescue effects of *Dc0^H2^*/+ on lifespan and its effects on further boosting the enhanced memory of *inc* mutants might be either a direct role of PKA, or a consequence of the restored sleep. (**F** and **G**) Representative images of whole-mount brain expressing mCD8-GFP in subtypes of mushroom body α/β_c_ (**F**) or α’/β’ (**G**) Kenyon cells in *wt*/control, *Dc0^H2^*/+ and *inc^1^*with or without *Dc0^H2^*/+ backgrounds. α/β_c_ neurons form discrete clusters in morphology, but not for α’/β’ neurons. Scale bar: 50 μm. (**H-L**) Statistics of the cluster number of α/β_c_ neurons (**H**), the number of lobes (**I** and **K**) and the volume of cell body and dendrites (**J** and **L**). n = 30-42 for **H** and **J**, 15-21 for **I**, and 32-36 for **L**. *p < 0.05; **p < 0.01; ***p < 0.001; ns, not significant. Error bars: mean ± SEM.

How might Inc interact with PKA in the regulation of memory? Both Inc and PKA are widely expressed in the fly brain but enriched in the mushroom body [35, 36, 73], a higher brain center that regulates both sleep and memory [19, 20, 79]. As shown above, *inc* mutants exhibit gross mushroom body structural defects (Figures 5C and 5F), and severe circuit defects in subtypes of mushroom body neurons, especially the α/β core (α/β_c_) and α’/β’ Kenyon cells have been previously reported in *inc* mutants [39]. In this regard, a simple and probably the most direct consideration is that the further enhanced memory of *inc* mutants by *Dc0* heterozygosity (Figures 7A-7D) might be a consequence of re-modulations of the mushroom body structure and circuits. To address this question, we expressed a GFP reporter in either α/β_c_ or α’/β’ Kenyon cells and evaluated the effects of Inc/PKA interaction in regulating the morphological properties of mushroom body subtype neurons. Consistent with previous findings [39], *inc* mutants showed an increased number of clusters, severely affected lobe structure, and increased volume of cell bodies and dendrites of α/β_c_ neurons (Figures 7F and 7H-7J). Interestingly, while the cluster number phenotype remained unaffected (Figure 7H), the severe lobe structural phenotype was even more frequently triggered by *Dc0* heterozygosity (Figure 7I), though cell body and dendritic volume phenotype was slightly suppressed (Figure 7J). In addition to α/β_c_ neurons, *inc* mutants almost did almost never show any phenotype in the number of lobes of α’/β’ neurons, but their cell body and dendritic volume was also increased (Figures 7G, 7K and 7L). Both lobe number and volume of cell bodies and dendrites of *inc* mutants were obviously enhanced by *Dc0* heterozygosity (Figures 7G, 7K and 7L).

Taken together, the further boosted memory of *inc* mutants with *Dc0* heterozygosity is unlikely achieved by restored mushroom body circuits. Instead, the stronger olfactory memory of *inc* mutants is likely contributed by overgrowth and abnormal arborization of these circuits. Elevated PKA signaling in *inc* mutants likely suppresses mushroom body Kenyon cell overgrowth to constrain excessive memory. In relation to the functions of sleep, the drastically restored sleep in *inc* mutants by *Dc0* heterozygosity might also contribute to physiological functions of sleep in promoting cognitive functions.

## Discussion

One of the major hypothesized functions of sleep is to restore brain plasticity from prior wake phases, and prepare the brain for another wake episode of learning and memory [6]. Consistently, learning and memory processes promotes sleep for memory consolidation [10, 42, 80, 81]. Furthermore, genetic and pharmacological sleep induction supports cognitive functions in health and disease [11, 42, 43, 82, 83]. Following this logic, interfering with sleep, e.g., in genetic scenarios of reduced sleep, should attenuate the efficacy of memory formation. In this study, we started from our perplexing observation that *Drosophila insomniac* mutants, characterized by severely decreased sleep, display robustly elevated scores of olfactory memories. We further found that *inc* loss-of-function (LOF) suppresses sleep through elevated PKA signaling in the mushroom body, which in turn constrains their excessive ability to form new memories, suggesting a “hyperplasticity” scenario on molecular, circuit and behavioral level. This hyperplasticity of *inc* mutants likely derives from increased mushroom body neurogenesis and subsequently overproduction of Kenyon cells with arborization defects, as already shown by a previous work [39]. In direct consequence, the setpoint between the need of sleep and strength of memory function seems to be changed/sensitized. Brains lacking *inc* seemingly counteract this hyperplasticity by elevating PKA signaling in the mushroom body Kenyon cells. While this elevated PKA signaling restricts excessive memory formation, it comes at the cost of reduced sleep levels and lifespan.

Our results thus suggest a specific and unexpected relation between excessive memory formation and sleep with a particular role of cAMP/PKA signaling. In addition, we offer a mechanistic interpretation for the sleep phenotypes of autism-associated *inc* mutants. Moreover, these results might also be relevant in the context of neurological diseases with developmental origins.

### *inc* LOF drives developmental behavioral hyperplasticity

It seems paradoxical that defective mushroom body morphology is coupled with stronger memory in *inc* mutants (Figure 7). Inc is active during a specific time window of *Drosophila* brain development [39]. The overproduction of the mushroom body Kenyon cells with structural defects in *inc* mutants suggests mushroom body to be the crucial neuron population from where Inc functions to regulate sleep, memory and mushroom body development [39]. Notably, we found that *Dc0^H2^*heterozygosity further exaggerates the mushroom body circuit structural phenotypes of *inc* mutants (Figures 7F-7L), but also boosts their memory function (Figures 7B and 7C) and increases their sleep level (Figures 4B and 7A). Thus, this overgrowth of mushroom body Kenyon cells might causally drive the cognitive hyperplasticity phenotypes of *inc* mutants.

We currently lack a mechanistic insight into how the mushroom body overgrowth connects to their setpoint change in generating excessive memory and dysregulating sleep. Loss of Inc generates more mushroom body Kenyon cells, which might enhance existing pathways or potentially establish novel circuits supporting stronger olfactory memory functions, and as a compensatory mechanism, elevated PKA signaling likely limits the mushroom body circuit phenotypes and subsequently suppresses the excessive memory.

### How might *inc* LOF promote memory?

As discussed above, *inc* LOF exhibited enhanced memory and downregulating PKA signaling in *inc* background further exacerbated this phenotype, suggesting an elevated PKA signaling in limiting the cognitive hyperplasticity of *inc* mutants. However, it is unknown how *inc* LOF triggers stronger olfactory memory at the first place. It is more likely that *inc* LOF promotes memory through an unknown signaling pathway in parallel to PKA signaling (Figure 7E).

Despite PKA signaling acting like a “brake” for excessive memory (Figure 7E), what might be the unknown signaling pathway directly mediating the memory-promoting effects of *inc* mutants? As an adaptor protein for Cullin-3 ubiquitin E3 ligase, Inc might promote the degradation of a direct target of Cullin-3 ligase, whose accumulation due to impaired ubiquitination and degradation might be responsible for the memory-promoting effects of *inc* mutants. In this regard, a potential target of Inc/Cullin-3 complex might be the dopaminergic signaling involved in both sleep and memory regulation, as it was previously shown to be important in mediating the sleep phenotypes of *inc* mutants [36]. Furthermore, given that *inc* mutants have profound mushroom body circuit phenotypes, it will be important in the future to screen for the interactions between Inc and other factors involved in regulating mushroom body development.

Interesting in this regard, mushroom body-specific expression of gain-of-function Rho kinase (ROCK or Rok) promotes memory formation [84], and triggers mushroom body structural defects [85], similar to *inc* mutants. While potential effects of ROCK gain-of-function on sleep have not been addressed, *inc* LOF might modulate Ras/Raf/ROCK signaling when promoting excessive memory.

### A potential hierarchical signaling cascade links Inc to PKA

What might be the exact action of Inc in regulating PKA signaling? The evolutionary conserved nature of Inc as an adaptor of Cullin-3 E3 ligase for ubiquitination of target proteins suggests that PKA Dc0 might be a direct target of Inc/Cullin-3. Indeed, *in vitro* studies suggest that PKA catalytic subunit is regulated through CHIP E3 ligase mediated ubiquitination and proteolysis [86]. However, there is no evidence for a role of Cullin-3 in PKA ubiquitination so far. Given the elevated PKA kinase activity in *inc* mutants (Figures 5C-5E) and that their sleep phenotypes can be substantially rescued by suppressing PKA kinase activity with the expression of an active form of its regulatory subunit (Figures 5G-5L), it is more plausible that Inc indirectly regulates PKA activity through cAMP/PKA signaling regulators, for example Neurofibromatosis-1 (Nf1) and/or adenylate cyclase Rutabaga (Rut) (Figures 3F-3H). Thus, it will be of great interest and importance in the future to explore the Inc/Cullin-3 ubiquitination targets driving the developmental behavioral hyperplasticity.

### A Yin and Yang of cAMP/PKA signaling in integrating sleep with memory

cAMP/PKA signaling was among the first identified pathways controlling learning and memory [87]. Importantly, cAMP/PKA signaling levels bidirectionally and negatively regulate sleep (Figure 3G) [32], consistent with our finding that the elevated PKA kinase activity of *inc* mutant directly drives their sleep deficits (Figures 5C-5L).

Notably, inducing sleep in cAMP adenylate cyclase mutant *rutabaga* (which should suffer reduced PKA activity) rescues their memory phenotypes [43]. In other words, lower cAMP/PKA signaling in *rutabaga* mutants might require excessive sleep for achieving a certain level of memory. Following this logic, as an alternative explanation for our data, it is also possible that the increased PKA activity levels of *inc* mutants might allow to generate robust memories at a lower sleep level, and restoring sleep by reducing PKA signaling might, as observed, further boost memory.

Interestingly in this regard, *Dc0* overexpression in the mushroom body suppresses memory at young age [33, 88]. Conversely, a mild reduction in the level of Dc0 by establishing *Dc0* heterozygosity also prominently suppresses age-associated memory decline, leaving memory unaffected at young age [33]. We here show that such genetic manipulation does not seem to provoke an obvious sleep phenotype in young *wt*/control animals (Figures 3A and 3D), while in *inc* background it specifically rescued the sleep defects (Figures 2A-2F and 3A-3E) and further boosted memory (Figures 7A-7D). Different from *rutabaga* mutants whose changes of PKA activity might be outside the physiological range, *Dc0* heterozygosity might mimic scenarios physiologically established under certain conditions, allowing for maintenance of longevity and memory [33].

Surprisingly, *Dc0* heterozygous animals are compromised in survival upon acute oxidative stress (Figure 6A), but they are ultimately longer-lived compared to *wt*/control (Figure 6B). Why does a reduction of PKA signaling promotes longevity but does not protect from oxidative stress? This might be explained by a differential regulation of downstream pathways by PKA in reacting to acute stress and chronic aging. Alternatively, PKA signaling may be differentially required for survival under acute oxidative stress and chronic stress driving aging. Indeed, acute H_2_O_2_ exposure directly activates PKA and subsequently phosphorylate substrate proteins *in vivo* [89], which might be critical for oxidative stress resistance. Interestingly, cAMP level, Dc0 protein level and PKA kinase activity in fly heads remain unchanged during early aging at age 20-day [33], while there is a significant reduction in Dc0 level of the 30-day fly wing neurons [90]. Thus, it might be possible that mushroom body cAMP/PKA signaling becomes adverse for longevity and memory during aging.

Thus, the functions of the cAMP/PKA signaling pathway regard the regulation of sleep, memory and survival might be dissectible in a spatial- and temporal-specific manner. The protected memory performance during aging and extended longevity of *Dc0^H2^* heterozygous animals are likely achieved at the sacrifice of resistance or tolerance to acute oxidative stress. The Yin and Yang is thus a plausible philosophical view on the complex cellular functions of cAMP/PKA signaling under distinct circumstances.

### Phosphorylation homeostasis as a mechanistic measure of sleep need for memory?

PKA as protein kinase has a broad range of phosphorylation targets, including essential synaptic proteins important for neurotransmission and synaptic plasticity [91–93]. In mice, AMP-activated protein kinase SIK3 was shown to be a target of PKA [94], whose phosphorylation by PKA inhibits its kinase function, consistent with the distinct roles of sleep regulation between SIK3 and PKA [95]. Specifically, SIK3 gain-of-function provokes a phosphorylation state of its substrates that promotes sleep [95], while PKA Dc0 gain-of-function suppresses sleep potentially by phosphorylating and counteracting SIK3 function and subsequently achieve a wake-promoting phosphorylation state [32]. In SIK3 *Sleepy* mice, while overall synaptic protein phosphorylation likely represents a signature of sleep need [95], whether and how their sleep behavior contributes to cognitive functions remain unclear. In *inc* mutants, it is possible that elevated PKA activity triggers a wake-promoting phosphorylation state representing the reduced levels of sleep need. Furthermore, *Dc0^H2^* heterozygosity likely renormalizes the phosphorylation state and subsequently the level of sleep of *inc* mutants, which further supports even stronger memory function (Figures 7B-7E). For both *inc* mutants and *Dc0* heterozygous animals, future study will employ quantitative proteomic and phosphoproteomic analyses to uncover their exact phosphorylation states in combination with sleep and memory screens, and also provide insights into age-associated memory decline.

### Neurodevelopmental hyperplasticity among neurological disorders

Collectively, by intensively studying the seemingly paradoxical *inc* short sleep mutants, we propose a cellular and behavioral hyperplasticity scenario originated from developmental overgrowth of mushroom body Kenyon cells, an essential brain center for sensory integration and sleep regulation. Our work identifies a molecular signaling cascade for programming a dynamic and plastic setpoint to tune cognitive hyperplasticity by measuring sleep amount at the sacrifice of longevity. While such behavioral hyperplasticity scenarios with sleep defects and cognitive disturbances are hallmarks of neurodevelopmental disorders like autism [21, 22], their potential etiology to these disorders remains obscure. As Inc functions as an adaptor for Cullin-3 E3 ligase-mediated ubiquitination, and Cullin-3 lesions have been associated with autism [40, 41], our current study may also provide mechanistic insights into the etiology of e.g., autistic patients.

## Materials and methods

### *Drosophila* stocks and maintenance

Flies were reared under standard laboratory conditions and raised on semi-defined medium (Bloomington recipe) under a 12/12 h light/dark cycle with 65% humidity at 25°C. 5- to 8-day-old male flies were used for all experiments except aversive olfactory memory experiments in which mixed populations of both sexes were used. All fly strains were backcrossed to *w^1118^* (*iso31*, BDSC#5905) background for at least six generations or retained in an already isogenized background. For the use of the two *inc* alleles in different experiments, due to the fact that *inc^2^* mutants are in general healthier than *inc^1^* (Figures 6D, 6F, 6N and 6P), *inc^2^* flies were more frequently used than *inc^1^*, especially for Pavlovian aversive olfactory conditioning experiments, in which large number of flies are required. However, due to the UAS cassette in the p-element of *inc^2^* mutants which drives *inc* expression when pairing with *Gal4* lines (Figure 1N) [35, 39], Gal4/UAS binary expression system was only used in *inc^1^* background (Figures 5C-5L and 7F-7L). Transgenic and mutant fly lines used in this study are listed below.

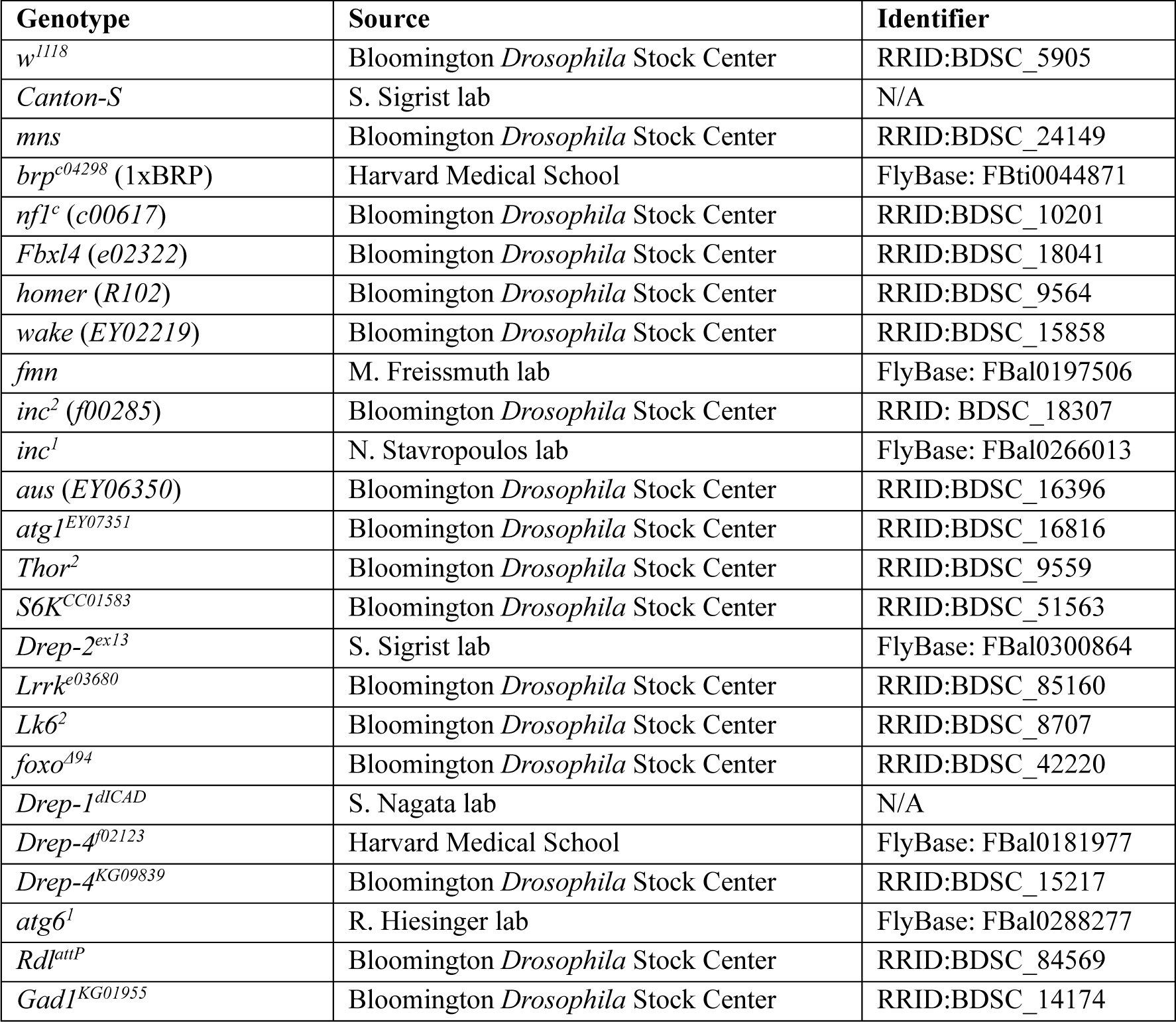
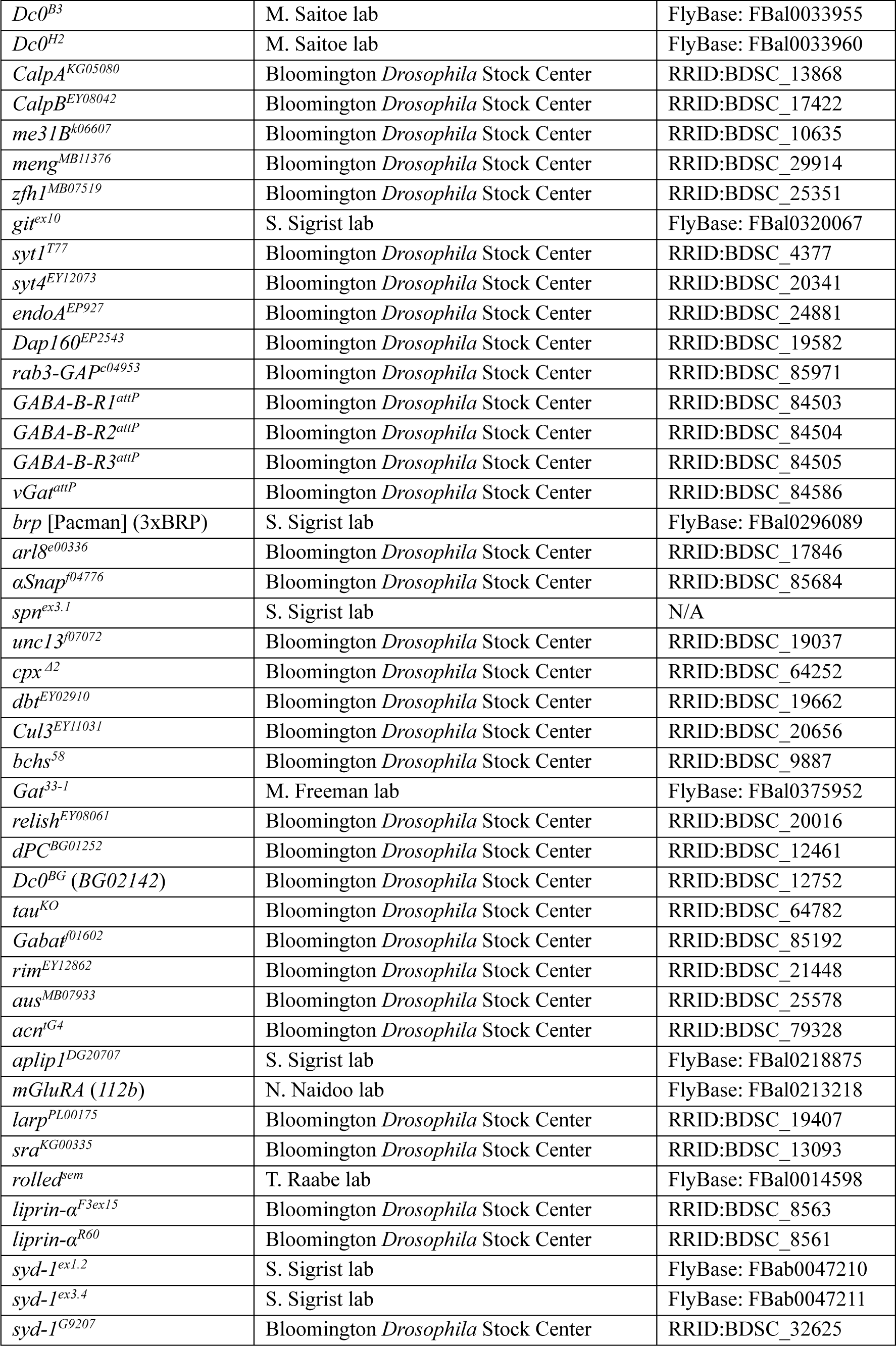
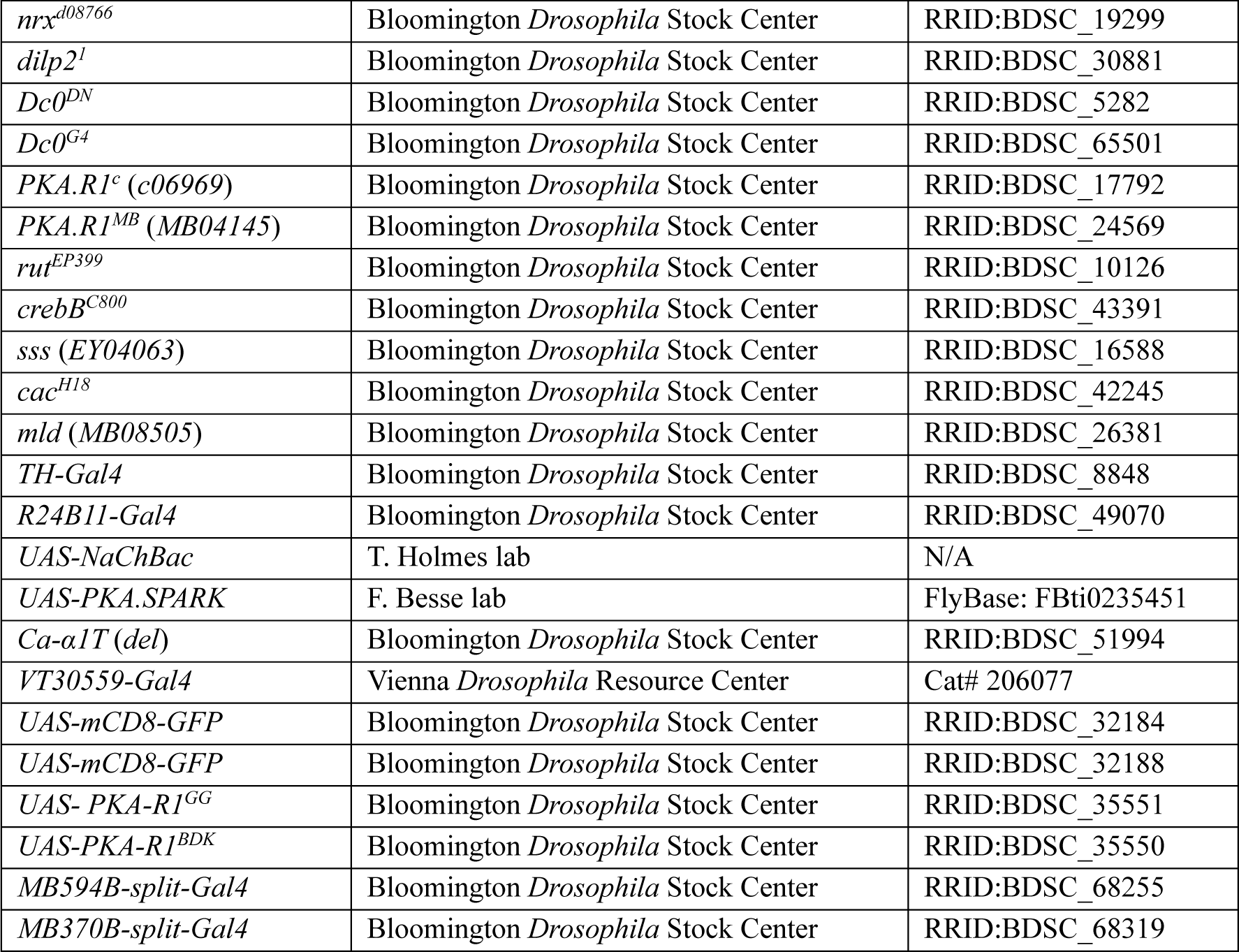

### Aversive olfactory learning and memory

Pavlovian aversive olfactory conditioning was performed as previous reported [11, 46]. Briefly, two aversive odors, 3-Octanol (OCT) and 4-methylcyclohexanol (MCH), served as behavioral cues (odors were diluted in mineral oil at a 1:100 ratio), and 120 V AC current electrical shocks were used as a behavioral reinforcer. Briefly, during one training session, around 100 naive flies were placed in a T-maze and exposed to the first odor (conditioned stimulus, CS+, MCH or OCT) paired with electric shock (unconditioned stimulus, US) for 60 s followed by 60 s rest, and then exposed to the second odor (CS-, OCT or MCH) without electric shocks for 60 s. During testing, these flies were simultaneously exposed to CS+ and CS- for 60 s to choose between the two odors. A reciprocal experiment, in which the second odor was paired with electric shocks as CS+, was performed. A performance index was calculated as the number of flies choosing CS- odor minus the number of flies choosing CS+ odor, divided by the total number of flies, and multiplied by 100 to get a percentage. The final performance index was the mean of the two performance indices from the reciprocal experiments.

STM was tested immediately after training, while 3h MTM was tested 3 h after training. As a component of 3h MTM, 3h ARM was separated from 3h ASM by cold shock. The trained flies received a cold shock on ice for 90 s, 2 h 30 min after training. 3h ARM was tested 30 min after the cold shock, and 3h ASM was calculated by subtracting 3h ARM from 3h MTM.

Odor avoidance was conducted similarly. Briefly, naive flies were placed in the choice position of the T-maze and allowed to choose between one of the two odors and air. The performance index was calculated as the number of flies choosing air minus the number of flies choosing the odor, divided by the total number of flies.

### Sleep measurements and genetic modifier sleep screen

Sleep experiments were performed exactly as previously reported [46]. Briefly, sleep of single male flies was measured by Trikinetics *Drosophila* Activity Monitors (DAM2) from Trikinetics Inc. (Waltham, MA) in 12/12 h light/dark cycle with 65% humidity at 25°C. 3∼4-day-old single male flies were loaded into Trikinetics glass tubes (5 mm inner diameter and 65 mm length) which have 5% sucrose and 2% agar in one side of the tube. Flies were entrained for at least 24 h and their locomotor activity was measured at 1 min interval. Due to entrainment to new environment, data from the first ∼24 h were excluded. A period of immobility without locomotor activity counts lasting for at least 5 min was determined as sleep [28]. Sleep from multiple days of recordings was analyzed and averaged using the Sleep and Circadian Analysis MATLAB Program (SCAMP) [96].

To carry out genetic modifier sleep screen for the X-linked *inc* mutants, we crossed female virgin *inc* mutants to either *wt* as control or isogenized autosomal candidate mutants to introduce heterozygosity of candidate mutations to *inc* male hemizygous mutants. For each measurement or biological replicate, differences between *inc* mutant with or without heterozygous mutations were calculated and then different measurements or replicates were pooled to generate Figure 2. At least two biological replicates were performed for each candidate mutant. As *inc* mutants, especially *inc^2^* mutants, show more profound sleep phenotypes during nighttime than during daytime, night P[wake] was specifically analyzed for the genetic modifier screen in Figure 2 so as to show major effects of different candidate heterozygous mutants on night sleep quality. For other P[wake] analysis, especially that *Dc0* heterozygosity has similar effects between day and night for *inc* mutants, daytime and nighttime were averaged for simplicity.

### Whole-mount brain immunostaining

Whole-mount brain immunostaining was performed similar to previous report [11, 46]. Brains of 5-day old male flies were used for all staining experiments. Flies were kept on ice for 1–2 min and then dissected in ice-cold Ringer’s solution (130 mM NaCl, 5 mM KCl, 2 mM MgCl_2_, 2 mM CaCl_2_, 5 mM HEPES, 36 mM Sucrose, pH = 7.3). Dissected brains were fixed immediately in 4% Paraformaldehyde (w/v) in PBS for 30 min on an orbital shaker at room temperature. Samples were washed in 0.7% Triton-X (v/v) in PBS (0.7 % PBST) for 20 min × 3 times, followed by blocking with 10% normal goat serum (v/v) in 0.7 % PBST for at least 2h at room temperature on shaker. After overnight antibody (rabbit anti-GFP conjugated with Alexa 488, 1:500; Nc82, 1:50) incubation at 4°C in darkness, brains were washed in 0.7% PBST for 30 min × 6 times at room temperature. Afterwards, for GFP Alexa 488 staining, brains were directly mounted on microscope slides in Vectashield and kept in a dark place at 4 °C before being scanned. For Nc82 staining, overnight secondary antibody (Cy5, 1:300) incubation at 4°C in darkness followed by 6 times washing in 0.7% PBST was performed before mounting.

### Confocal microscopy and image analysis

Whole-mount adult brain samples were imaged on a Leica TCS SP8 confocal microscope (Leica Microsystems), and images were obtained using the Leica LCS AF software (Leica Microsystems). For analyzing PKA SPARK intensity and puncta, the mushroom body lobes were imaged using a 63× 1.40 NA oil objective with a voxel size of 0.2405 µm × 0.2405 µm × 1 µm at a speed of 400 Hz. For morphology analysis, mushroom body lobes and calyx were imaged using a 40× 1.30 NA oil objective with a voxel size of 0.3788 µm × 0.3788 µm × 1 µm. All the parameters were kept constant for the same set of experiments. Image stacks were processed and analyzed with Fiji (https://fiji.sc/).

### Mushroom body calyx and cell body volume analysis

The quantification of calyx volume was performed semi-automatically via a custom-written Fiji Macro, with steps briefly as follows. Selected stacks containing regions of cell bodies and dendritic processes of the calyx were converted into binary images after background subtraction, image smoothing, and thresholding processes, and then the volumes were measured using the Plugin Voxel counter.

### PKA SPARK analysis

For PKA SPARK experiments, imaging stacks of mushroom body lobes were converted to two-dimensional images using max Z-projection. The average gray pixel value of the PKA SPARK was measured within a ROI of mushroom body lobes segmented automatically after background subtraction, image filtering, and thresholding processes. The SPARK puncta were detected using “Find Maxima”.

### Oxidative stress survival assay

Male flies were sorted into a population of 45–50 flies per vial at 2 days post-eclosion. One day after sorting, the flies were transferred into a vial containing 4 layers of filter paper soaked with 1.4 ml 5% sucrose solution with 2% H_2_O_2_. Flies were transferred into new vials with freshly prepared filter papers and solution every other day. The numbers of deaths were counted and recorded daily at the same time of the day. Comparison of survival curves was performed with GraphPad Prism using Log-rank analysis.

### Longevity

Longevity experiments were carried out exactly as previously reported [11]. Briefly, male flies were sorted into a population of approximately 20 to 25 flies per vial/replicate at age 2-day-old. To reduce the variability in lifespan, a few different cohorts of flies were used for each experiment. The longevity flies were regularly transferred onto fresh food every second day or over the weekend and the number of dead flies in each vial was recorded at each time of transfer until the death of the last fly of a replicate. Comparison of longevity curves was performed with GraphPad Prism using Log-rank analysis.

### Statistics

Statistics was performed as previously described [46]. GraphPad Prism 6 was used to perform most statistics and figures were generated using both Adobe Illustrator and Prism. Student’s t test was used for comparison between two groups and one-way ANOVA with Tukey’s post hoc tests were used for multiple comparisons between multiple groups (≥3). For mushroom body lobe number analysis (Figures 7I and 7K), Kruskal-Wallis test with Dunn’s multiple comparisons was used.

## Lead contact and materials availability

Further information and requests for resources should be directed to and will be fulfilled by Sheng Huang as well as Stephan J. Sigrist. This study did not generate new unique reagents.

## Acknowledgement

We thank N. Stavropoulos, M. Saitoe, F. Besse, R. Hiesinger, N. Naidoo, M. Freeman, M. Freissmuth, S. Nagata, Bloomington *Drosophila* Stock Center (BDSC), Harvard Medical School, and Vienna *Drosophila* Resource Center for fly lines. This work was supported by grants from European research council (ERC) advanced grant and the Deutsche Forschungsgemeinschaft (DFG; German Research Foundation) to S.J.S. (SFB1315 TP A08 and FOR2705 TP05). S.H. was supported by the Leibniz SAW SyMetAge and Charité NeuroCure.

## Disclosure and competing interests statement

The authors declare no competing interests.

## Author contributions

Conceptualization, S.H.; Investigation, S.H., C.P., Z.Z., C.B.B., O.T., and D.T.; Funding acquisition and supervision: S.H., and S.J.S.; Writing, S.H. with inputs from other authors.

